# The MscS channel from *Corynebacterium glutamicum* uses a non-canonical gating mechanism

**DOI:** 10.64898/2026.09.02.748947

**Authors:** Yoshitaka Nakayama, Giorgos Hiotis, Thomas Walz

## Abstract

The function and gating mechanism of *Escherichia coli* mechanosensitive channel of small conductance (*Ec*MscS) is well understood, but it is unknown whether MscS homologs in other bacteria function the same way. Here, we show that the MscS homolog from *Corynebacterium glutamicum* (*Cg*MscS) opens at a much higher membrane tension than *Ec*MscS but has otherwise similar functional characteristics. *Cg*MscS is also structurally similar to *Ec*MscS but features an extended transmembrane (TM) helix 2 that forms salt bridges with the cytoplasmic cage. Compared to the closed conformation, the TM1-2 sensor paddles in the inactivated conformation are rotated but not tilted, and, unlike *Ec*MscS, there is no change in the associated pocket lipids, thus establishing a gating mechanism distinct from *Ec*MscS that is not based on the “lipids-move-first” model. Because the TM2 extension is conserved in Actinobacteria but not other bacterial phyla, our findings suggest that MscS homologs have lineage-specific gating mechanisms.

## INTRODUCTION

Mechanosensitive (MS) channels are membrane proteins that convert mechanical force into electrical signals and play central roles in diverse biological processes, ranging from osmoregulation, to hearing and the sense of touch, to blood pressure regulation (1). The first MS channels were discovered in bacteria and were named MS channels of large and small conductance, MscL and MscS, respectively (2,3). The structure and function of *Escherichia coli* MscS (*Ec*MscS) have been studied extensively. The channel is closed in a relaxed membrane and opens when the membrane tension reaches 5.5 - 6.2 mN/m, forming a channel with a conductance of ∼1.0 nS (4,5). Channel gating is based on the “force-from-lipids” mechanism, which posits that the channel opens in response to changes in the transmembrane (TM) pressure profile, which can be caused, for example, by changes in membrane tension or membrane curvature (6,7). *Ec*MscS has fast gating kinetics, so that it can respond quickly to hypoosmotic shocks, and it shows little gating hysteresis, closing at a similar membrane tension as when it opens (8). In addition, *Ec*MscS features sub-conducting states and can desensitize and inactivate (9,10). Several mutations have been identified that change the channel characteristics of *Ec*MscS. Most notably, mutation of Gly113, which forms a kink in the pore-lining helix, to bulkier residues reduces both inactivation of MscS and its sensitivity to an increase in membrane tension (11,12). Substitutions of Ala106 with residues that have larger side chains also make the channel harder to open (13).

The structure of *Ec*MscS was first determined by X-ray crystallography (14). The channel is a homoheptamer, with each protomer contributing three TM helices, TM1-3. TM1 and TM2 at the periphery of the channel form a “sensor paddle” that responds to membrane tension, whereas TM3 is divided by a kink at Gly113 into the pore-lining helix TM3a and the amphipathic helix TM3b that connects the pore to the cytoplasmic domain. The cytoplasmic domain is a cage-like structure that is formed by the membrane-proximal β subdomains and the membrane-distal αβ domains, which, at their interfaces, form lateral openings, referred to as fenestrations. The first crystal structure, which did not allow modeling of side chains in the pore region, was initially thought to represent the channel in an open conformation, mainly due to the narrowest pore diameter still appearing to be ∼7 Å (14–16). Later, this structure was recognized as a non-conductive state, because the hydrophobic Leu105 and Leu109 residues constrict the pore diameter to ∼5 Å and form a vapor-lock mechanism that de-wets the pore (17). A subsequent study using the A106V loss-of-function mutant revealed a drastically different conformation, in which the sensor paddles were more tilted and the TM3a pore helices were rotated clockwise, resulting in an increase in the pore diameter to ∼13 Å, thus revealing the conformation of *Ec*MscS in an open state (18). The same open conformation was later seen for wild-type (WT) *Ec*MscS in a single-particle cryogenic electron microscopy (cryo-EM) structure obtained after extended incubation with detergent to delipidate the channel (19). Recent cryo-EM structures were determined of MscS reconstituted into membrane-scaffold protein (MSP)-based lipid nanodiscs, which revealed that numerous lipids associate with the channel (20–22). By modifying the lipid membrane, i.e., by using very short-chain lipids to create a very thin membrane and by removing lipids from the nanodisc to mimic membrane tension, it was also possible to visualize *Ec*MscS in other functional states, which were interpreted to represent the sub-conducting and desensitized states, respectively (22), and provided a structural basis for the “lipids-move-first” model, which proposed that pocket lipids had to dissociate before the channel can transition into the open conformation (23).

MscS is the founding member of the superfamily of MscS-like channels, with members being found in bacteria (24–27), archaea (28), fungi (29), plants (30–32), and parasites (33). MscS-like channels are structurally and functionally diverse (34,35). They can be smaller than MscS, missing part of the cytoplasmic domain and/or some TM helices, as for example *Nematocida displodere* MscS2 (36) and *Trypanosoma cruzi* MscS (37), or much larger, featuring, for example, additional TM helices and periplasmic domains, as in the case of *E. coli* MscK (38). Even though most MscS-like channels contain a core structure resembling MscS, structural studies revealed that they use gating mechanisms that differ from the lipids-move-first mechanism used by MscS (23,39). For example, gating of YnaI entails only very small conformational changes that mostly involve the straightening of the pore-lining helix (40), while gating of MscK involves flattening and expansion of the entire TM domain and a conformational rearrangement of all 77 TM helices (38). The functional characteristics of MscS-like channels are equally diverse, as illustrated by YnaI, which, unlike MscS, only has two functional states, closed and open, opens at near-lytic membrane tension, has slow gating kinetics and pronounced gating hysteresis (41,42).

The MscS-like channels MscCG (also known as NCgl1221) and MscCG2 expressed in *Corynebacterium glutamicum* are of high industrial importance (43). *C. glutamicum* is used for the production of glutamate, the “umami” taste enhancer, and its two MscCG channels provide the major pathways for glutamate excretion (44,45). Historically, the NCgl1221 gene was found to be mutated in *C. glutamicum* strains that spontaneously excrete glutamate, and the protein encoded by the gene was initially proposed to be a glutamate exporter. However, due to the similarity of its N-terminal region with *Ec*MscS, the gene product was later renamed to mechanosensitive channel of *Corynebacterium glutamicum* (MscCG) (46). Deletion of the MscCG gene reduces glutamate excretion by ∼80%, but not completely, which prompted the prediction that a secondary glutamate exporter must exist. Eventually, the MscCG2 gene was found in the genome of most industrial but not all *C. glutamicum* strains (45). First electrophysiological characterizations using *C. glutamicum* giant spheroplasts of the industrial strain ATCC13689 revealed that MscCG and MscCG2 differ from each other in terms of mechanosensitivity and conductance (47). MscCG opens at a membrane tension of ∼5.5 mN/m and forms a pore with the rather small conductance of ∼340 pS, while MscCG2 opens at a near-lytic tension of ∼12 mN/m and then forms a pore with a conductance of ∼1.0 nS (47). These different channel characteristics suggest that MscCG and MscCG2 have distinct activation mechanisms that allow them to respond to different degrees of membrane tension.

*C. glutamicum* usually only excretes glutamate when MscCG opens in response to increased membrane tension, which can be induced by biotin-limiting conditions (44). However, the A111V mutation in the pore of MscCG results in constitutive glutamate excretion (44). Electrophysiological analysis of the A111V mutant showed that it has a smaller activation threshold and increased gating hysteresis compared to WT MscCG, so that it is easier to open and then remains open for a longer time (48). Overexpression of MscCG2 carrying the A151V mutation, which corresponds to the A111V mutation in MscCG, also causes constitutive glutamate excretion, but overexpression of the two mutant MscCG channels has very different effects. While overexpression of A111V MscCG severely impairs cell growth (49), overexpression of A151V MscCG2 further increases the yield of glutamate in the medium under biotin-limiting conditions (45), which prompted an interest in ion-channel engineering to further improve the efficiency of MscCG2-mediated glutamate release and thus glutamate production (50,51).

The structure of MscCG2, predicted using the Deep-learning Iterative Threading ASSEmbly Refinement (D-I-TASSER) program (52), looked very similar to that of *Ec*MscS, with the notable exception that helix TM2 has a cytoplasmic extension that reaches the cytoplasmic cage (53). Based on this prediction, the A151V mutation would be located in the kink of pore helix TM3, which is similar to mutation of Gly113 in *Ec*MscS and may thus have a similar effect on channel function. However, it has not yet been experimentally tested whether MscCG2 is gated in the same way as *Ec*MscS. This question needs to be addressed, because MscCG2 functions in a very different environment than *Ec*MscS. *C. glutamicum* is a Gram-positive bacterium, which, unlike Gram-negative *E. coli*, has a thick and waxy cell wall with a mycolic acid layer that provides additional protection to hypoosmotic shock and the membrane of *C. glutamicum*, unlike that of *E. coli*, consists predominantly of phosphatidylglycerol and cardiolipin and is thus highly negatively charged (54,55).

In this study, we combine cryo-EM and patch-clamp electrophysiology to determine how MscCG2 senses membrane tension and gates in the context of its native membrane. We show that MscCG2 preserves the canonical MscS architecture, and so we will refer to it as *C. glutamicum* MscS (*Cg*MscS), but it contains an extended TM2 helix that links the membrane-embedded sensor paddle to the cytoplasmic cage. Despite its high activation threshold, *Cg*MscS exhibits clear inactivation. Structural analysis further shows that *Cg*MscS gating involves a rotation of the sensor paddles and twisting of the pore-lining TM3a helices, but does not seem to involve tilting of the sensor paddles, which is a critical part of *Ec*MscS gating. These results establish a distinct gating mechanism that is based on the cytoplasmic extension of TM2 coupling the sensor paddle to the cytoplasmic cage. Comparative sequence analysis reveals that the TM2 extension is specific to Actinobacteria, indicating that the distinct gating mechanism may be a common feature of MscS homologs from Actinobacteria.

## RESULTS

### Anionic lipids stabilize *Cg*MscS gating

The channel properties of *Ec*MscS (5,11,56) and MscCG (46,47,57) have already been characterized, but much less is known about the functional characteristics of *Cg*MscS. We therefore expressed WT *Cg*MscS in *E. coli* strain MJF612, in which the predominant MS channels (MscL, MscS, YbdG and MscK) have been deleted (58), and prepared giant spheroplasts for analysis by patch-clamp electrophysiology. Application of negative pressure to the patch membrane elicited highly flickery currents that prevented reliable single-channel measurements (**Fig. 1a**). We previously recorded endogenous MS channel currents from giant spheroplasts prepared from the industrial *C. glutamicum* strain ATCC13869, which exhibited more stable channel openings and a high activation threshold, comparable to that of *C. glutamicum* MscL (*Cg*MscL) (47) (**Fig. 1b**). To test whether it is the native *C. glutamicum* membrane lipids that stabilize *Cg*MscS gating, we used a previously developed fused vesicle system (59). We expressed *Cg*MscS in a *C. glutamicum* strain ATCC13032-derived strain lacking MscCG, *Cg*MscL, and *Cg*MscS, and prepared membrane vesicles. These membrane vesicles were fused with soy polar lipid (SPL) liposomes at a 1:1 ratio, and these fusion vesicles were used for patch-clamp experiments. Upon application of negative pressure to patches excised from the fusion vesicles, stepwise *Cg*MscS currents suitable for single-channel analysis were observed (**Fig. 1c**). These currents were less flickery than those recorded from *E. coli* membranes, which contain ∼70% zwitterionic phosphatidylethanolamine (PE) lipids, but remained more flickery than those observed in native *C. glutamicum* membranes, which contain ∼70% negatively charged phosphatidylglycerol (PG) lipids.

**Fig. 1.**
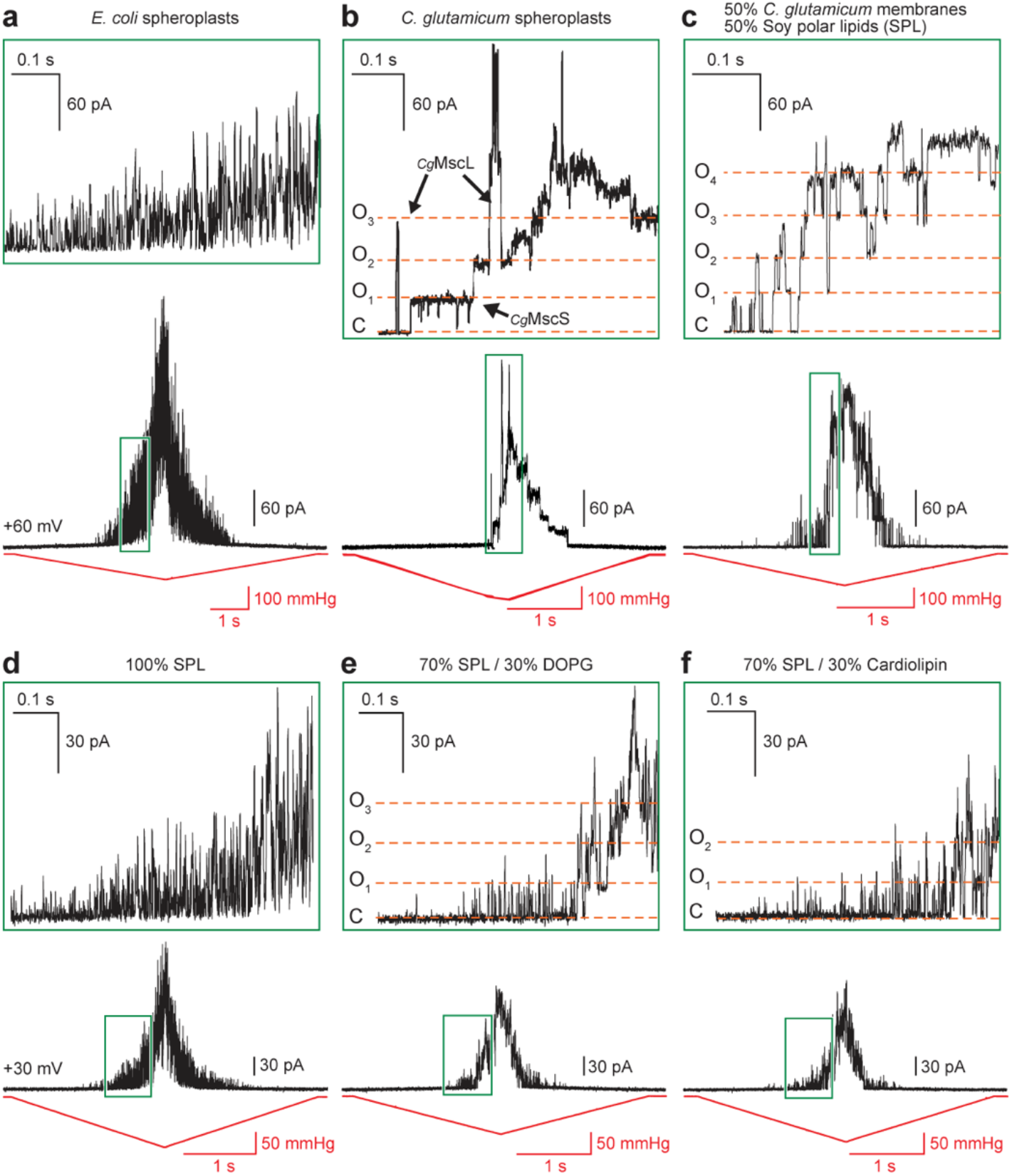
Anionic lipids stabilize the open state of *Cg*MscS. **a-c** Representative traces of currents in response to ramp-pressure stimuli recorded from giant spheroplasts prepared from *E. coli* expressing *Cg*MscS (a), giant spheroplasts prepared from *C. glutamicum* expressing *Cg*MscS (adapted from Nakayama *et al.*, *Sci Rep* 2018) (b), and fusion vesicles obtained by fusing membrane vesicles prepared from *Cg*MscS-expressing *C. glutamicum* with soy polar lipids (SPL) liposomes (c). **d-f** Representative traces of currents in response to ramp-pressure stimuli recorded from *Cg*MscS proteoliposomes reconstituted with 100% SPL (d), 70% SPL and 30% DOPG (e), and 70% SPL and 30% cardiolipin (f). The top panels are zoomed-in views of the regions indicated by the boxes in the bottom panels to show single-channel current levels. Red line: pressure; black line: current; horizontal orange dashed lines: levels when all *Cg*MscS channels are closed (C) and when one, two, three or four *Cg*MscS channels are open (O1 – O4).

To identify which lipids of the *C. glutamicum* membrane contribute to the stabilization of channel gating, we first reconstituted purified *Cg*MscS into liposomes with SPL, which contain ∼50% zwitterionic phosphatidylcholine (PC) lipids. Under these conditions, elicited currents were similarly flickery as those observed in *E. coli* giant spheroplasts (**Fig. 1d**), indicating that *Cg*MscS gating is unstable in membranes rich in zwitterionic lipids. *C. glutamicum* membranes are rich in anionic lipids, primarily PG lipids, so we tested whether anionic lipids can stabilize *Cg*MscS gating. Doping of the SPL liposomes with either 30% dioeleoyl PG (DOPG) or cardiolipin improved the stepwise gating behavior of reconstituted *Cg*MscS (**Fig. 1e, f**). These findings indicate that stabilization of *Cg*MscS gating does not depend on a specific lipid species but instead reflects a general effect of anionic lipids.

### *Cg*MscS has *Ec*MscS-like channel characteristics but a much higher activation threshold

We used the fusion vesicles described above to further examine the channel properties of *Cg*MscS and compare them with those of *Ec*MscS and the glutamate-release channel MscCG. *Ec*MscS is characterized by rapid activation and strong inactivation (11,60), whereas MscCG displays slower activation kinetics and little or no inactivation (57). A pressure-ramp protocol elicited currents from all three channels, confirming channel activity under the same recording conditions. *Cg*MscS activated at ∼75 mmHg and closed at ∼70 mmHg during pressure release (**Fig. 2a**, top). Quantitation showed that *Cg*MscS required a significantly higher activation pressure of 73.1 ± 2.6 mmHg (mean ± standard error of the mean [s.e.m.]; *n* = 7) (**Fig. 2d**) than either *Ec*MscS with 34.6 ± 4.5 mmHg (mean ± s.e.m.; *n* = 5) (**Fig. 2b**, top, and **2d**) or MscCG with 35.6 ± 4.5 mmHg (mean ± s.e.m.; *n* = 7) (**Fig. 2c**, top, and **2d**), consistent with previous recordings in *C. glutamicum* giant spheroplasts (47). Using a step-pressure protocol, we observed rapid activation of *Cg*MscS followed by gradual current decay during sustained pressure (**Fig. 2a**, bottom). *Ec*MscS showed similar rapid activation followed by gradual current decay under sustained pressure (**Fig. 2b**, bottom), consistent with its known inactivation behavior. In contrast, MscCG exhibited slower activation kinetics and sustained currents with no detectable inactivation under sustained pressure (**Fig. 2c**, bottom).

**Fig. 2.**
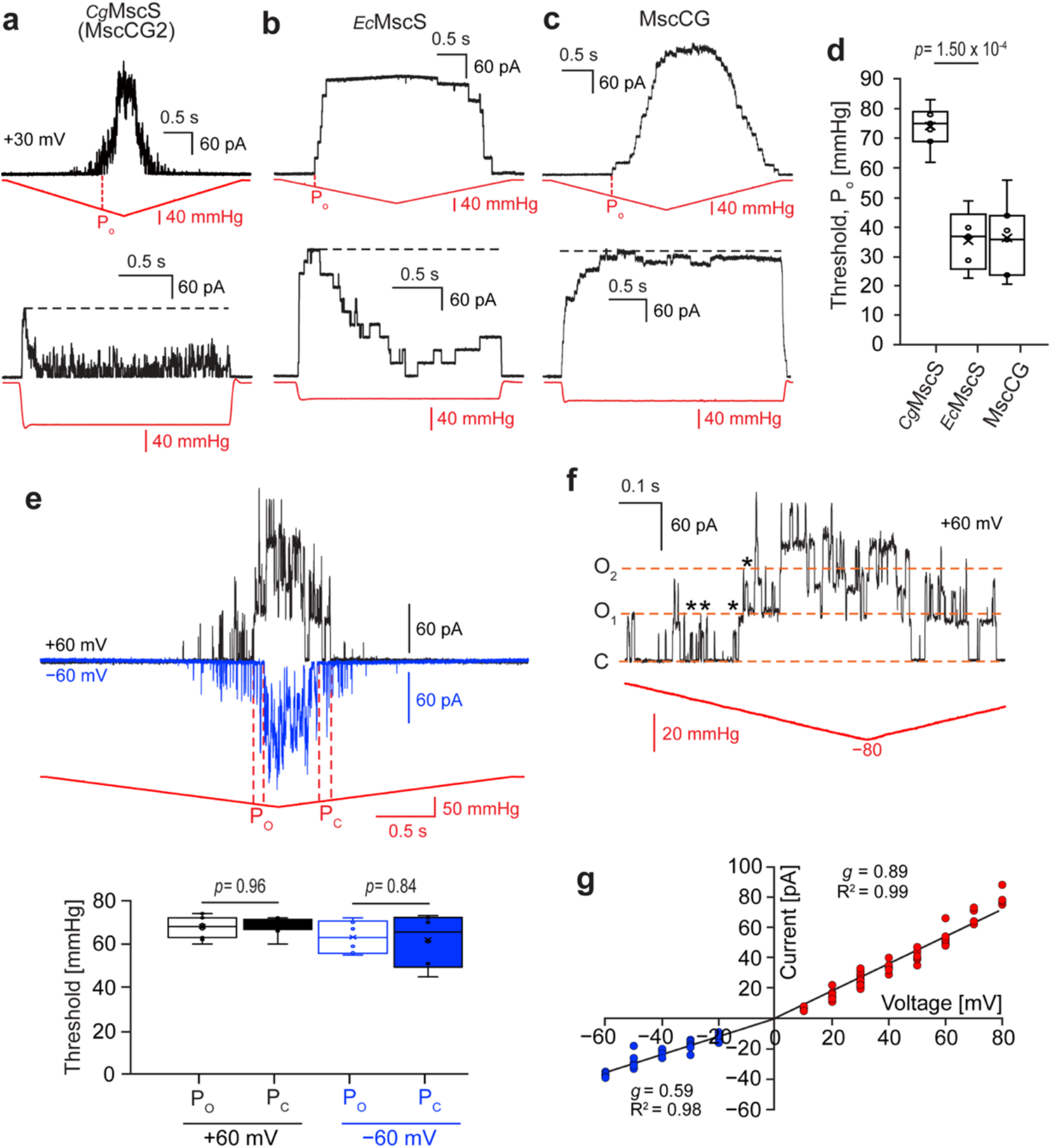
Channel characteristics of *Cg*MscS. **a-c** Representative traces of currents in response to ramp (top panels) and step (bottom panels) pressure stimuli. Panel a shows current traces recorded from fusion vesicles formed from *Cg*MscS-containing *C. glutamicum* membrane vesicles and SPL liposomes (n = 21). The step-pressure recording shows rapid *Cg*MscS activation followed by current decay during sustained pressure. Panel b shows current traces recorded from fusion vesicles formed from *Ec*MscS-containing *E. coli* membrane vesicles and SPL liposomes (n = 5). Panel c shows current traces recorded from fusion vesicles formed from MscCG-containing *C. glutamicum* membrane vesicles and SPL liposomes (n = 7). The step-pressure recording shows slower activation and sustained currents with little or no inactivation. Red line: pressure; black line: current; P_o_ in top panels: first full channel opening; horizontal black dashed lines in bottom panels: peak current levels. **d** Comparison of activation thresholds, measured as the pressure of the first full channel opening, for *Cg*MscS (n = 7), *Ec*MscS (n = 5) and MscCG (n = 7). Statistical analysis: two-sided Welch’s unequal-variance *t*-test. Box plot shows median (centre line), interquartile range (box) and full data range (whiskers). **e** Top panel: Representative traces of currents in response to ramp-pressure stimuli recorded from *Cg*MscS-containing fusion vesicles at voltages of +60 mV and −60 mV (n = 6). Red line: pressure; black/blue line: current; P_o_: first full channel opening; P_c_: last channel closing. Bottom panel: The opening and closing thresholds show no statistically significant differences, indicating that the channels have no gating hysteresis at both positive and negative voltages. Statistical analysis: two-sided Welch’s unequal-variance *t*-test. Box plot shows median (centre line), interquartile range (box) and full data range (whiskers). **f** Single-channel conductance analysis of WT *Cg*MscS. Currents were recorded from *Cg*MscS-containing fusion vesicles at +60 mV (n = 11). Red line: pressure; black line: current; horizontal orange dashed lines: levels when all *Cg*MscS channels are closed (C) and when one or two *Cg*MscS channels are open (O1, O2), asterisks: subconductive levels. **g** Current–voltage relationship for *Cg*MscS. Currents were recorded from *Cg*MscS-containing fusion vesicles at voltages ranging from −60 mV to +80 mV (more than three independent patch membranes per voltage). The slope conductances (*g*) and coefficients of determination (R²) were calculated from independent linear fits to the measurements obtained with positive (red) and negative (blue) voltages.

Unlike *Ec*MscS (61), *Cg*MscS showed no obvious gating hysteresis at either positive or negative voltages (**Fig. 2e**, top). To quanify the voltage dependency of the gating hysteresis, the pressures at which the first full channel opening (Po) and the last channel closing (Pc) occurred were measured at voltages of +60 mV (black) and −60 mV (blue). The Po values at +60 mV and −60 mV were 67.2 ± 1.6 mmHg (mean ± s.e.m.; *n* = 10) and 62.2 ± 3.0 mmHg (mean ± s.e.m.; *n* = 6), respectively, whereas the Pc values were 67.3 ± 1.2 mmHg (mean ± s.e.m.; *n* = 10) and 61.0 ± 4.8 mmHg (mean ± s.e.m.; *n* = 6), respectively (**Fig. 2e**, bottom). These results indicate that *Cg*MscS shows little if any gating hysteresis and that its activation and closing thresholds are not strongly voltage-dependent.

*Cg*MscS displayed multiple subconductance levels (**Fig. 2f**), making it impossible to reliably estimate the single-channel conductance of fully open *Cg*MscS. Therefore, to minimize the variation in the conductance values used to establish the current–voltage relationship, currents were measured at multiple voltages with more than three independent recordings (**Fig. 2g**). From the slopes of the graph at positive and negative voltages, we determined conducatnces of 890 pS and 590 pS, respectively, indicating that *Cg*MscS has an overall conductance comparable to that of *Ec*MscS (∼1 nS) and exhibits weak current rectification. Together, these results show that *Cg*MscS has *Ec*MscS-like conductance and inactivation but requires a substantially higher membrane tension for activation.

### *Cg*MscS features a cytoplasmic extension of TM2, lipid-filled pockets, and a flexible αβ subdomain

The unusual gating features that distinguish *Cg*MscS from both *Ec*MscS and MscCG prompted us to determine a structure for this MscS homolog. We expressed WT *Cg*MscS natively in *C. glutamicum* and purified it in the detergent dodecyl maltoside (DDM). Because PG lipids stabilized the gating of *Cg*MscS, we reconstituted the channel with DOPG into nanodiscs formed with the membrane-scaffold protein (MSP) MSP1E3D1 and used cryo-EM to determine a density map at an overall resolution of 2.9 Å, which allowed us to build an atomic model for most of the protein (**Fig. 3a**, left, and **Fig. S1**). *Cg*MscS subunits have the conserved MscS fold and assemble into a homoheptamer (**Fig. 3a**, left) as orginally established for archetypal *Ec*MscS (**Fig. 3a**, right). The N-terminal sequence forms an amphipathic helix and the TM domain of *Cg*MscS adopts a splayed conformation, in which the TM helices are close together at the periplasmic side of the membrane but then tilt outwards relative to the channel axis and are thus further apart on the cytoplasmic side. This structure resembles the conformation seen for *Ec*MscS in the closed state. Consistent with this finding, analysis with the program HOLE (62) revealed that the narrowest region of the pore at residue Phe149 has a diameter of 4.4 Å, confirming that this conformation represents a non-conductive, closed state (**Fig. 3b**).

**Fig. 3.**
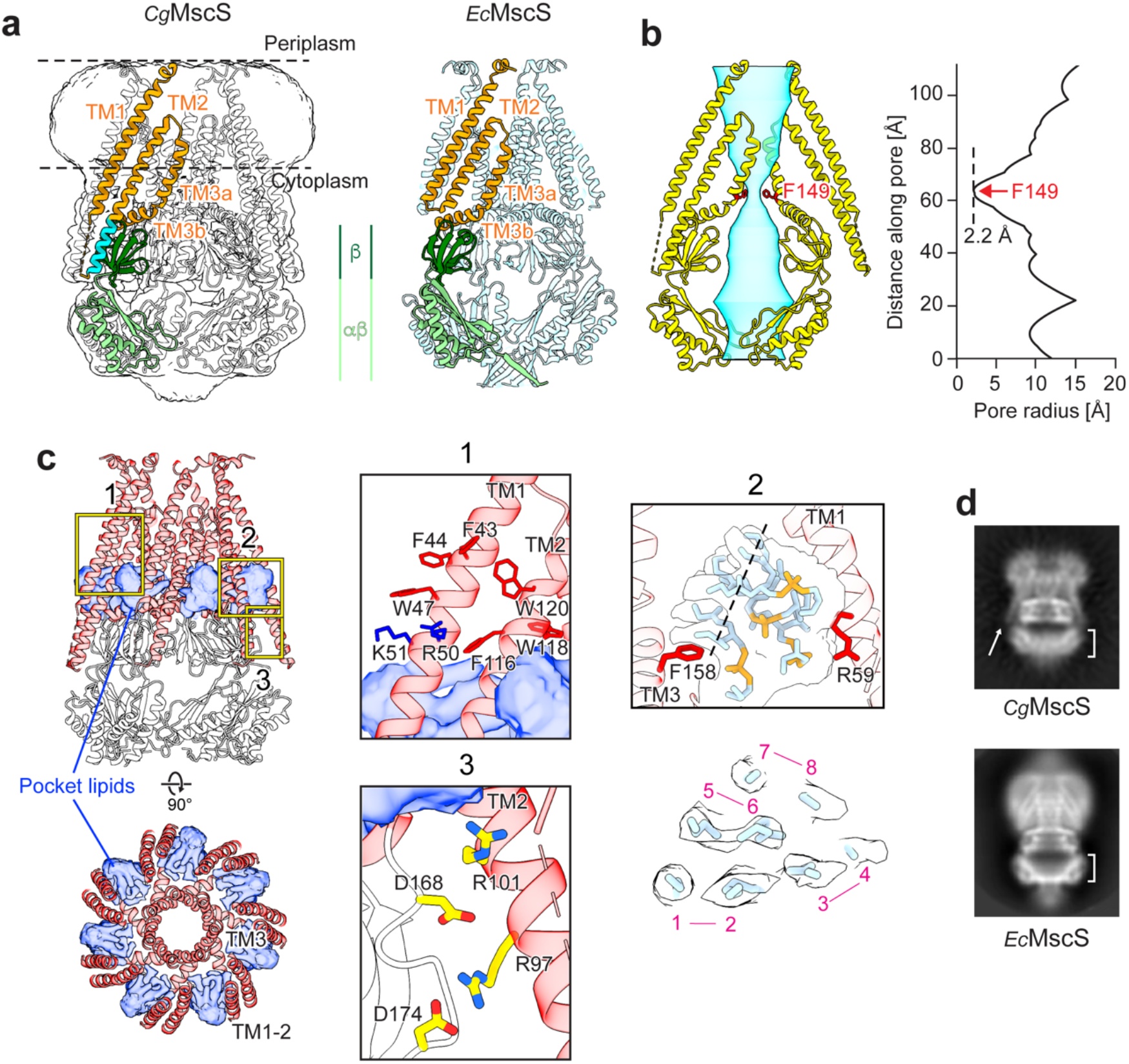
Cryo-EM structure of WT *Cg*MscS in DOPG nanodiscs. **a** Cryo-EM structures of *Cg*MscS in DOPG nanodiscs (left) and *Ec*MscS in DOPC nanodics (right; PDB: 6VYK) (22). The *Cg*MscS density map is shown as transparent white surface, and the models for the *Cg*MscS and *Ec*MscS heptamers are shown in white and light blue, respectively. One protomer in each channel is color-coded: orange: TM1-2 (sensor paddle), TM3a (pore-lining helix) and TM3b; green: cytoplasmic cage (β subdomain in dark green; αβ subdomain in light green); cyan: TM2 extension unique to *Cg*MscS. The horizontal dashed lines indicate the membrane boundary predicted by the Orientations of Proteins in Membranes database (https://opm.phar.umich.edu). **b** The volume representation (left panel) and linear profile of the pore radius (right panel) generated with HOLE (62) show that the pore in *Cg*MscS is closed and that the constriction site is formed by the aromatic Phe149 residue. The vertical dashed line shows the pore radius at the constriction site. **c** Views perpendicular (top panel) and parallel to the membrane (bottom panel) of the *Cg*MscS heptamer shown in red (TM helices) and white (cytoplasmic cage) ribbon representation. Densities occupying the hydrophobic extramembranous cavities that likely represent pocket lipids are shown as blue transparent surfaces. Box 1 shows a zoomed-in view of aromatic residues (Trp and Phe; red) and positively charged residues (Arg and Lys; blue) near the cytoplasmic membrane boundary that may anchor the position of TM1 and TM2. Box 2 shows a zoomed-in view of the four lipids modeled into the pocket density, and the panel below shows a section through the pocket density at the position indicated in Box 2, revealing density for the eight acyl chains. Box 3 shows a zoomed-in view of the putative electrostatic interactions between arginine residues in the TM2 extension and aspartate residues in the β subdomain that link the sensor paddle to the cytoplasmic cage. **d** 2D-class averages showing side views of *Cg*MscS (top panel) and *Ec*MscS (bottom panel, from Zhang et al., Nature, 2022). The arrow indicates the extended TM2 in *Cg*MscS and the brackets show the αβ subdomain that is well-defined in *Ec*MscS but smeared out in *Cg*MscS.

*Cg*MscS differs from *Ec*MscS in four ways. First, while the pore constriction in *Ec*MscS is formed by two leucine residues, Leu105 and Leu109, the pore constriction in *Cg*MscS is formed by the single Phe149 residue (**Fig. 3b**). Second, TM2 of *Cg*MscS features a cytoplasmic extension that reaches the β subdomain of the cytoplasmic cage, suggesting a different mechanical coupling between the TM1-2 sensor paddle and the cytoplasmic cage (**Fig. 3a**). Third, unlike *Ec*MscS, *Cg*MscS has several aromatic tryptophan and phenylalanine and positively charged arginine and lysine residues near the cytoplasmic membrane boundary in TM1 and TM2 that likely anchor the vertical position of the TM1-2 paddle in the membrane (**Fig. 3c**, inset 1). Fourth, while the cytoplasmic cage of *Ec*MscS is very rigid and therefore always best resolved in cryo-EM density maps of this channel, only the β subdomain is well-resolved in our map of *Cg*MscS while the αβ subdomain has lower resolution (**Fig. S1d**), indicating that this domain is not rigidly connected to the β subdomain (see below).

The *Cg*MscS map did not show density for gatekeeper or pore lipids seen for *Ec*MscS in the closed conformation (21,22). However, like *Ec*MscS, the TM domain of *Cg*MscS does contain hydrophobic cavities that are occupied by density not accounted for by the protein (**Fig. 3c** and **S4a**, bottom panels). The pocket density shows eight tubular features, indicating that the pockets are filled by four lipids, with their putative headgroups located near Arg59 and their acyl chains located near Phe158, consistent with electrostatic and hydrophobic interactions of *Cg*MscS with pocket lipids, respectively (**Fig. 3c**, inset 2). Like in *Ec*MscS, TM2 forms one side of these pockets but then it extends further into the cytoplasm. This cytoplasmic extension of TM2 contains positively charged residues Arg97 and Arg101, which are close to negatively charged residues Asp168 and Asp174 in the β subdomain, thus creating potential electrostatic interactions that link the sensor paddle to the cytoplasmic cage (**Fig. 3c**, inset 3).

Density for the loop between TM1 and TM2 and for the C-terminal end that in *Ec*MscS forms a cap closing the cytoplasmic cage was missing in the map of *Cg*MscS, and the αβ subdomain was more poorly resolved than the β subdomain (**Fig. S1d**), indicating structural variability in these regions. This can indeed already be seen in two-dimensional (2D) class averages. While class averages of *Ec*MscS show well-defined features for the entire cytoplasmic domain, class averages of *Cg*MscS only show well-defined features for the β but not the αβ subdomain (**Fig. 3d**). To examine the conformational variability of the αβ subdomain, we subjected it to three-dimensional (3D) variability analysis using a focused mask, which revealed a breathing motion, showing that the αβ subdomain can expand and contract (**Movie S1**). We then performed 3D classification focused on the αβ subdomain, which yielded classes, in which the αβ subdomain showed different degrees of radial expansion **(Fig. S1h**). We selected three classes, representing the most contracted, an intermediate, and the most expanded conformation, and built atomic models of the αβ subdomain for each class, which could be modeled confidently up to residue Pro313 (**Fig. S2**).

Overlay of the three models reveals the breathing motion, showing the peripheral helices of the αβ subdomain progressively moving radially outwards from the most contracted to the intermediate to the most expanded conformation (**Fig. 4a**). HOLE analysis shows that this motion does not affect the pore constriction at Phe149, while the radius profiles within the αβ subdomain differ substantially (**Fig. 4b**). In particular, the diameter of the axial opening of the cage, measured at the position of the last modeled residue, Pro313, is ∼20 Å in the most contracted conformation, but increases to ∼30 Å in both the intermediate and most expanded conformations (**Fig. 4b** and **Fig. 4c**, middle panels). However, the expansion is not uniform across the αβ subdomain, being most pronounced at the axial opening of the cage and barly noticeable close to the β subdomain. As a result, the lateral fenestrations formed at the interface between the β and αβ subdomains are very similar in the three conformations (**Fig. 4c**, bottom panels), so that ion access to the channel through the lateral fenestrations should not be affected by the breathing motion of the αβ subdomains. However, while the lateral fenestrations provide the only access to the channel in *Ec*MscS, in which the axial opening to the cage is closed by a cap structure, the axial opening in *Cg*MscS is large and is unobstructed, making it unclear how important the lateral fenetartions are for providing ions access to the channel.

**Fig. 4.**
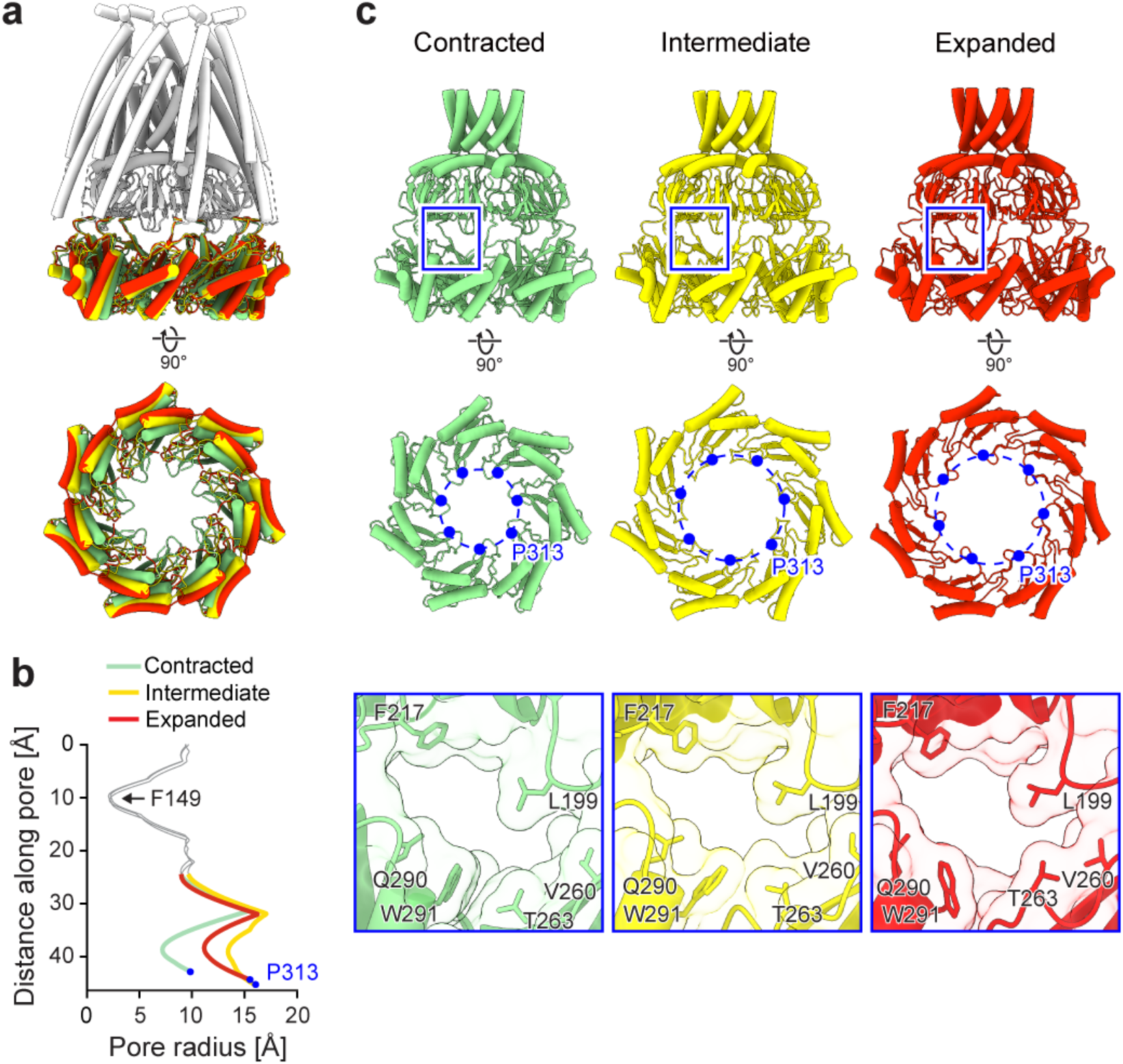
Expansion/contraction of the αβ subdomain of WT *Cg*MscS. **a** Overlay of the most contracted (light green), intermediate (yellow), and most expanded (red) conformations of the αβ subdomain. Views perpendicular (top panels) and parallel to the membrane (bottom panels) are shown. **b** HOLE radius profiles in the region of TM3, the β subdomain, and the αβ subdomain (residues 134–313) of *Cg*MscS with the αβ subdomain in the three different conformations. The portions of the profiles corresponding to TM3 and the β subdomain are shown in gray, whereas those corresponding to the αβ subdomain in the most contracted, intermediate and most expanded conformation are shown in light green, yellow and red, respectively. The constriction at Phe149, indicated by the arrow, is essentially unchanged between the three conformations. The blue filled circles mark the pore radius at the level of the last modeled residue, Pro313, illustrating that the radius of the axial opening in the cage is smaller in the most contracted conformation. **c** Comparison of the αβ subdomain and the lateral fenestrations in the most contracted (light green), intermediate (yellow), and most expanded (red) conformations. Views are shown parallel to the membrane for the cytoplasmic cage and TM3 (top panels) and perpendicular to the membrane from the cytoplasmic side for the αβ subdomain (middle panels). The blue filled circles mark the positions of Pro313, and the blue dashed circles indicate the diameter of the ring formed by the Pro313 residues. The bottom panels show close-up views of the lateral fenestrations indicated by the boxes in the top panels, illustrating that the openings are similar.

### The pocket lipids form a static structural element of *Cg*MscS

Patch-clamp experiments revealed that *Cg*MscS gating is stable in PG but not PC lipids, which we reasoned could be caused by different lipid species occupying the extraembranous pockets. We therefore determined the structure of *Cg*MscS in DOPC nanodiscs (**Fig. S3**). The resulting model was identical to that obtained with DOPG nanodiscs with a root-mean-square deviation between the backbone atoms of 0.489 Å (**Fig. S4b**, top panel). The density representing the pocket lipids was also essentially the same (**Fig. S4a, b**, middle and bottom panels), indicating that the same number of lipids occupy the pockets in a similar manner. Finally, we also determined the structure of *Cg*MscS purified in 0.05% DDM (**Fig. S5**) as well as after an overnight incubation in a ten-times higher DDM concentration (**Fig. S6**), an approach that was previously used to delipidate *Ec*MscS (19) and YnaI (63), which yielded structures of these channels in the open conformation. Both structures in detergent showed *Cg*MscS in the closed conformation (**Fig. S4c, d**, top panels) and, importantly, showed the same density for the pocket lipids as previously seen in DOPG and DOPC nanodiscs (**Fig. S4c, d**, middle and bottom panels). These results show that unlike the pocket lipids in *Ec*MscS, which form an important functional and dynamic part in its channel gating, the pocket lipids in *Cg*MscS are more of a structural and static element of the channel, ressembling those in MscS-like channel YnaI (42).

### The A151V *Cg*MscS mutant opens at a lower pressure and does not inactivate

The A151V mutation of *Cg*MscS causes constitutive glutamate excretion (44,45). This residue corresponds to Leu111 in *Ec*MscS in the kink region of TM3 (**Fig. 5a**), and mutations in this region alter *Ec*MscS gating (56). We therefore used *Cg*MscS fusion vesicles to test whether the A151V mutation similarly affects the gating of *Cg*MscS. In patch-clamp recordings using a pressure-ramp protocol, A151V *Cg*MscS activated at a significantly lower pressure than WT *Cg*MscS, namely 20.2 ± 3.7 mmHg (mean ± s.e.m.; *n* = 5) as compared to 73.1 ± 2.6 mmHg (mean ± s.e.m.; *n* = 7) (**Fig. 5b**, top panel, and **5c**), and the thresholds for the first full channel opening, P_o_, and the last channel closing, P_c_, were almost the same, suggesting that the A151V mutant does not have gating hysteresis (**Fig. 5b**, top panel). Furthermore, unlike for WT *Cg*MscS, the current of the A151V mutant did not decay under sustained pressure (**Fig. 5b**, bottom panel), and the channels spontaneously opened even after complete pressure release (which did not occur before pressure application) (**Fig. 5b**, top panel). However, single-channel analysis showed altered pore-conductance properties, with reduced single-channel conductance compared to WT *Cg*MscS and multiple subconductance levels (**Fig. 5d**). We therefore calculated the conductance from the slope of the current–voltage relationship, as described for WT *Cg*MscS, which was 500 pS at positive voltages and 450 pS at negative voltages (**Fig. 5e**). Together, these results show that the A151V mutation converts *Cg*MscS into a low-threshold, non-inactivating channel with altered pore conductance properties, with the spontaneous opening of the mutant channel providing a possible explanation for the constitutive glutamate excretion observed for *C. glutamicum* expressing A151V *Cg*MscS (44,45).

**Fig. 5.**
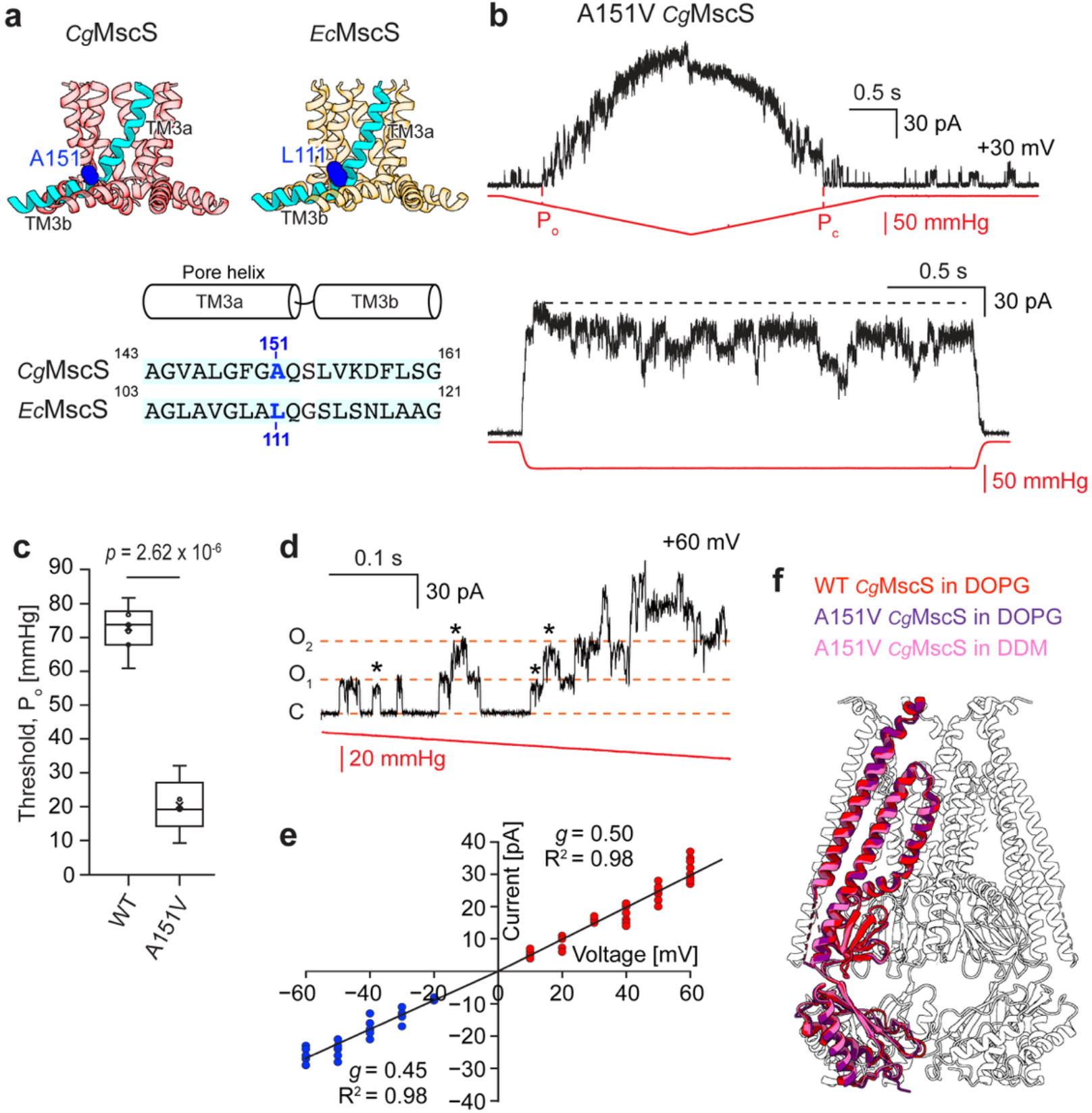
Functional and structural characterization of A151V *Cg*MscS. **a** Position of the A151V mutation in the kink region of the pore-lining TM3a helix of *Cg*MscS (transparent red, top left panel) and the corresponding residue L111 in *Ec*MscS (transparent orange, top right panel). One TM3 helix is shown in cyan, and residues A151 in *Cg*MscS and L111 in *Ec*MscS are shown as blue spheres. The bottom panel shows a sequence alignment of the TM3 segments surrounding the kink region in *Cg*MscS and *Ec*MscS, with the corresponding residues highlighted in blue. **b** Patch-clamp recordings of A151V *Cg*MscS in fusion vesicles in response to ramp (top panel) and step (bottom panel) pressure stimuli (n = 18, each). Red line: pressure; black line: current; P_o_: first full channel opening; P_c_: last channel closing; horizontal black dashed line: peak current level. **c** Comparison of activation thresholds, measured as the pressure of the first full channel opening, for WT *Cg*MscS (n = 7) and A151V *Cg*MscS (n = 5). Statistical analysis: two-sided Welch’s unequal-variance t-test. Box plot shows median (center line), interquartile range (box), and full data range (whiskers). **d** Single-channel conductance analysis of A151V *Cg*MscS. Currents were recorded from *Cg*MscS-containing fusion vesicles at +60 mV (n = 18). Red line: pressure; black line: current; horizontal orange dashed lines: levels when all *Cg*MscS channels are closed (C) and when one or two *Cg*MscS channels are open (O1, O2), asterisks: subconductive levels. **e** Current–voltage relationship for A151V *Cg*MscS. Currents were recorded from A151V *Cg*MscS-containing fusion vesicles at voltages ranging from −60 mV to +60 mV (more than three independent patch membranes per voltage). The slope conductances (*g*) and coefficients of determination (R²) were calculated from independent linear fits to the measurements obtained with positive (red) and negative (blue) voltages. **f** Overlay of the cryo-EM structures of WT *Cg*MscS in DOPG nanodiscs (red), A151V *Cg*MscS in DOPG nanodiscs (purple), and A151V *Cg*MscS in DDM (pink), showing that the A151V mutant remains in a closed conformation.

The reduced activation threshold and stabilized conductive state of the A151V mutant raised the possibility that the mutation also stabilizes *Cg*MscS in the open conformation. We therefore determined cryo-EM structures of A151V *Cg*MscS both in DOPG nanodiscs (**Fig. S7**), to provide a native membrane environment, and in DDM (**Fig. S8**), because structures of *Ec*MscS in the open conformation could only be determined in detergent (22). In both environments, A151V *Cg*MscS adopted essentially the same closed conformation seen for WT *Cg*MscS in DOPG (**Fig. 5f**).

### β-cyclodextrin (βCD) does not allow visualization of *Cg*MscS in an open conformation

βCD-mediated lipid extraction, which mimics membrane tension, is a general approach to activate MS channels (64) and made it possible to determine the structure of *Ec*MscS in a desensitized conformation (22). We therefore tested whether we could use βCD treatment of *Cg*MscS-containing nanodiscs to obtain a structure of the channel in an open conformation. However, cryo-EM analysis of WT *Cg*MscS in DOPG nanodiscs after a 16-h incubation with 100 mM βCD yielded a structure indistinguishable from the conformation of *Cg*MscS in DOPG nanodiscs before βCD treatement (**Fig. S9** and **S10a**). We therefore tested whether it would be possible to visualize an open conformation if we used the A151V mutant, which has a lower activation threshold. However, cryo-EM analysis of A151V *Cg*MscS in DOPG nanodiscs after βCD incubation also showed the channel in the closed conformation (**Fig. S11** and **S10a**).

To verify that βCD-mediated lipid removal actually can activate *Cg*MscS, we performed patch-clamp experiments on *Cg*MscS-containing fusion vesicles. As a control, we first applied a pressure ramp to *Ec*MscS channels, which showed the presence of active channels, and then added βCD during the same recording from the same patch membrane, which revealed progressive, stepwise channel openings before the membrane finally ruptured (**Fig. S10b**). We then performed the same experiment with A151V *Cg*MscS. Application of a pressure ramp confirmed the presence of active channels, and subsequent addition of βCD treatment resulted in stepwise, although flickery single-channel openings before the membrane ruptured (**Fig. S10c**). These observations demonstrate that βCD can increase membrane tension sufficiently to activate A151V *Cg*MscS channels, and that it is a limitation of the nanodisc system that prevents visualization of the open conformation of *Cg*MscS in βCD-treated nanodiscs, as was previously the case for YnaI (42).

### A146V *Cg*MscS reveals the structure of an inactivated channel

The structure of the A106V *Ec*MscS mutant in the open conformation showed that gating of this channel involves a rotation and tilting of the TM1-2 sensor-paddle helices and a straightening of the pore-lining TM3a helices (18) (**Fig. 6a**). In the closed conformation, the TM3a helices are tilted and Ala106 packs closely against Gly108 in the neighboring TM3a (**Fig. 6b**, top panel). Mutation of Ala106 to valine interferes with this packing interaction and leads to a straightening of the TM3a helices and opening of the pore (**Fig. 6b**, middle panel). The *Cg*MscS residue that corresponds to *Ec*MscS residue Ala106 is Ala146, and it forms a similar diagonal packing interaction with Gly148 in the neighboring TM3a, in the closed conformation (**Fig. 6b**, bottom panel). We therefore introduced the corresponding A146V mutation in *Cg*MscS with the expectation that it would allow us to visualize *Cg*MscS in the open conformation. However, after cryo-EM analysis of A146V *Cg*MscS in DOPG nanodiscs (**Fig. S12**), the overall structure looked very similar to that of WT *Cg*MscS, including the breathing motions of the αβ subdomain (**Fig. S13**), and did not show the expected tilt of the TM1-2 paddles (**Fig. 6c**). However, in the A146V mutant, the TM1-2 paddle is rotated by ∼52°, which corresponds to a transition of the position of one subunit in the heptameric assembly to that of the neighboring subunit. As in *Ec*MscS, this rotation changes the organization of the monomers from a domain-swapped configuration in the closed state, in which the TM1-2 paddle of one subunit aligns with the β subdomain of the neighboring subunit, to an aligned configuration, in which the TM1-2 paddle of a subunit aligns with its own β subdomain (**Fig. 6a, c**). In both channels, the rotation of the TM1-2 paddle results in a twisting of pore-lining helix TM3a, which changes from a tilted to a more membrane-perpendicular orientation. However, in contrast to *Ec*MscS, in which the conformational change includes a tilt of the TM1-2 paddle that results in a thinning of the TM domain, the TM1-2 paddle in *Cg*MscS does not tilt and the thickness of the TM domain remains unchanged. Furthermore, the density in the extramembraneous pockets of A146V *Cg*MscS appeared comparable to that in WT *Cg*MscS in DOPG nanodiscs (**Fig. S14**), so that the conformational change is unlikely to be caused by delipidation of the pockets.

**Fig. 6.**
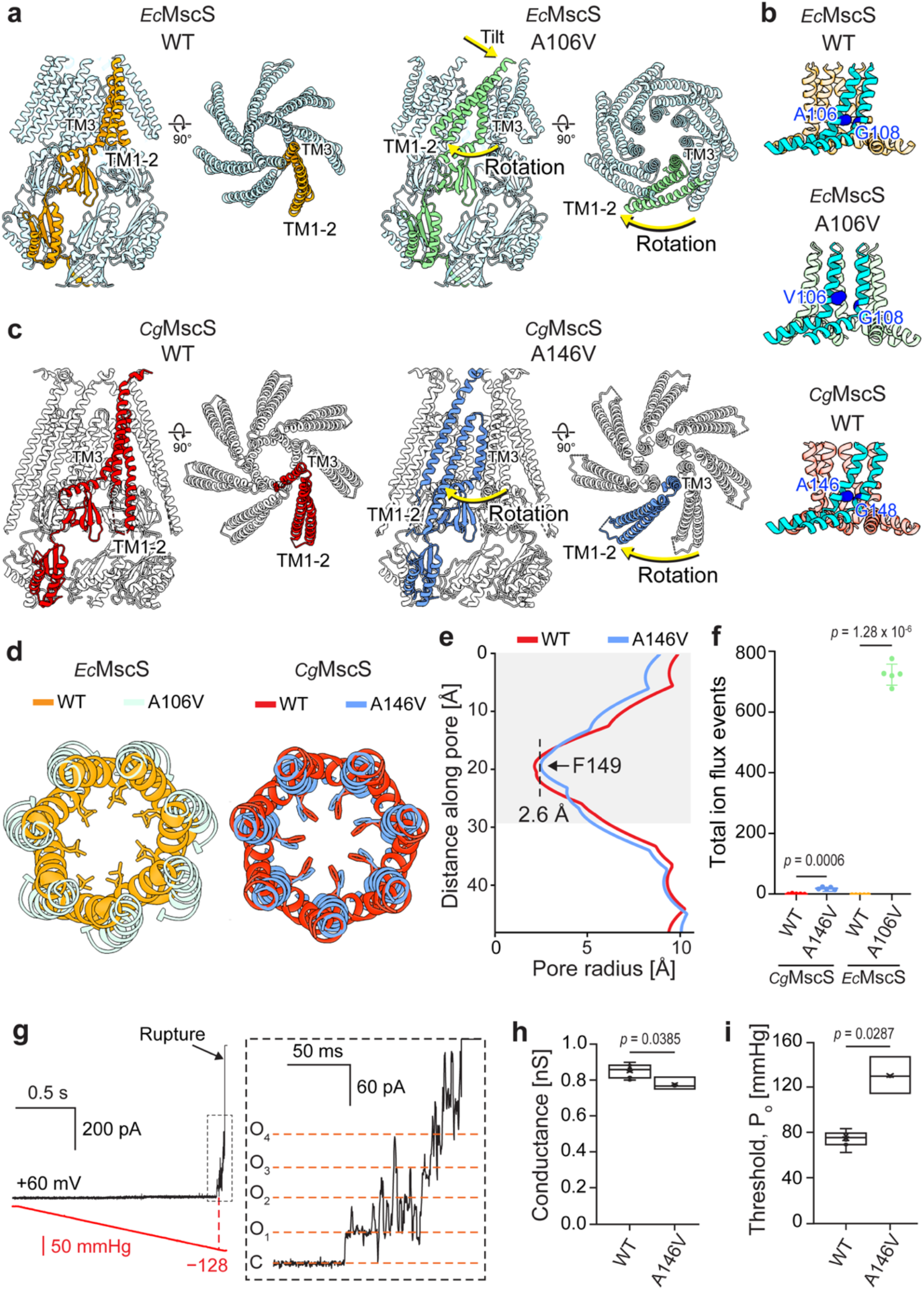
Structural and functional characterization of A146V *Cg*MscS. **a** Cryo-EM structures of WT *Ec*MscS in the closed conformation (left; PDB: 2OAU; one protomer in orange) (14) and A106V *Ec*MscS in open conformation (right; PDB: 2VV5; one protomer in green) (18), showing that the TM1-2 paddle rotates and tilts. **b** Intersubunit packing interactions between adjacent TM3a helices in WT *Ec*MscS in the closed state (top), A106V *Ec*MscS in the open state (middle), and WT *Cg*MscS in the closed state (bottom). Two neighboring protomers are shown in cyan, and glycine and alanine residues involved in helical packing interactions are shown as blue spheres. **c** Cryo-EM structures of WT *Cg*MscS in the closed conformation (left, one protomer in red) and A146V *Cg*MscS in the inactivated conformation (right; one protomer in blue), showing that the TM1-2 paddle only undergoes a rotation but no tilt. **d** Arrangement of the TM3a helices in closed WT *Ec*MscS (orange) and open A106V *Ec*MscS (light green) (left), and in closed WT *Cg*MscS (red) and inactivated A146V *Cg*MscS (blue) (right). The side chains of Leu105 and Leu109 in *Ec*MscS and of Phe149 in *Cg*MscS, which form the pore constrictions, are shown. **e** Pore radius profiles generated with HOLE for WT (red) and A146V *Cg*MscS (blue). The light gray box indicates the pore region lined by the TM3a helices. **f** Total ion-flux events observed in 200-ns molecular-dynamics simulations for WT and mutant *Ec*MscS and *Cg*MscS. Data plotted as mean **±** SD (n = 5). Statistical analysis: two-sided Welch’s unequal-variance *t*-test. **g** Currents were recorded from A146V *Cg*MscS-containing fusion vesicles in response to increasing negative pressure (n = 3). Red line: pressure; black line: current; vertical red dashed line: first full channel opening. The right panel shows a zoomed-in view of the region indicated by a box in the left panel to resolve single-channel conductance levels. Horizontal dashed lines: levels when all channels are closed (C) and when one, two, three or four channels are open (O1 – O4). **h, i** Comparison of channel conductance (h) for WT (n = 10) and A146V *Cg*MscS (n = 3) and of activation thresholds, measured as the pressure of the first full channel opening (i), for WT (n = 7) and A146V *Cg*MscS (n = 3). Statistical analysis: two-sided Welch’s unequal-variance t-test. Box plots show median (center line), interquartile range (box), and full data range (whiskers).

The conformational change in *Ec*MscS causes a dilation of the pore formed by the TM3a helices and the removal of Leu105 and Leu109, which form the constriction site, from the ion-conducting pathway, resulting in an open pore (**Fig. 6d**, left panel). In contrast, the conformational change in *Cg*MscS does not cause a significant pore dilation, and despite the ∼52° rotation, Phe149, which forms the pore constriction, ends up almost in the same position (**Fig. 6d**, right panel). Indeed, analysis with HOLE shows that the diameter of the constriction site in A146V *Cg*MscS is 5.2 Å and thus only slightly wider than the constriction-site diameter of 4.4 Å in WT *Cg*MscS (**Fig. 6e**).

The constriction-site diameter of A146V *Cg*MscS of 5.2 Å is much smaller than that of *Ec*MscS A106V of ∼13 Å (18), which suggested that this *Cg*MscS conformation may not be conductive. To test this possibility, we performed all-atom molecular-dynamics simulations. While WT *Ec*MscS was non-conductive with 0 ions permeating the pore of the channel over 200 ns of simulation under a constant electric field of +200 mV, A106V *Ec*MscS showed high ion conductivity with 723.4 ± 15.6 ions permeating the pore (**Fig. 6f**). In contrast, both WT and A146V *Cg*MscS showed no physiologically meaningful ion conduction with 2.0 ± 0.5 and 18.2 ± 2.1 ions permeating the pores, respectively (**Fig. 6f**).

Taken together, the conformation adopted by A146V *Cg*MscS, in which the channel has undergone a conformational change but is not ion-conductive, likely represents the inactivated state of this channel.

### The A146V mutation in *Cg*MscS is functionally equivalent to the A106V mutation in *Ec*MscS

Although the A146V mutation in *Cg*MscS corresponds to the A106V mutation in *Ec*MscS, the cryo-EM structures of these two mutants revealed different functional states, prompting us to use patch-clamp electrophysiology to characterize A146V *Cg*MscS and to establish whether the two mutants are functionally equivalent. Patch-clamp recordings showed that A146V *Cg*MscS forms functional MS channels; however, this mutant required substantially higher membrane tension for activation than WT *Cg*MscS, approaching the membrane lytic tension. Using a pressure-ramp protocol, the first A146V *Cg*MscS channel opened at a negative pressure of 128 mmHg, immediately before membrane rupture (**Fig. 6g**). This behavior is consistent with the previous characterization of A106V *Ec*MscS as a loss-of-function variant (18). Furthermore, a single-channel conductance analysis showed that the conductance of A146V *Cg*MscS is 778 ± 35 pS (mean ± s.e.m.; *n* = 3), comparable to that of WT channels of 848 ± 12 pS (mean ± s.e.m.; n = 10) (**Fig. 6h**). However, the activation threshold of A146V *Cg*MscS was measured to be 128.7 ± 9.2 mmHg (mean ± s.e.m.; *n* = 3), which is significantly higher than that of WT *Cg*MscS of 73.1 ± 2.6 mmHg (mean ± s.e.m.; *n* = 7) **(Fig. 6i**). These results demonstrate that the A146V mutation in *Cg*MscS is indeed functionally equivalent to the A106V mutation in *Ec*MscS.

### The TM2–β-subdomain coupling is highly conserved in MscS homologs from Gram-positive bacteria

Our structures show that *Cg*MscS uses a gating mechanism that is distinct from the canonical gating mechanism that was established for the MscS homolog from the Gram-negative model bacterium *E. coli*. We thus wondered whether this distinct gating mechanism is unique to *Cg*MscS or is a common trait of MscS homologs from Gram-positive bacteria. The most distinct structural element in *Cg*MscS is the cytoplasmic extension of the TM2 that contains arginine residues interacting with aspartate residues in the β-subdomain in the cytoplasmic cage (**Fig. 3c**, inset3), which is likely a key structural feature underlying the distict gating mechanism of *Cg*MscS. To determine whether electrostatic coupling between TM2 and the β subdomain is a conserved feature, we identified representative three-TM-helix MscS homologs from Gram-negative bacteria (6,505; based on similarity with *Ec*MscS), the low-GC Gram-positive Firmicutes (1,414; based on similarity with *Bacillus subtilis* MscS [*Bs*MscS]), and the high-GC Gram-positive Actinobacteria (2,031; based on similarity with *Cg*MscS), and mapped the sequence conservation within each group onto the structures of *Ec*MscS, *Bs*MscS, and *Cg*MscS, respectively. Conservation mapping and sequence-logo analyses revealed highly conserved arginine residues in the TM2 extension and highly conserved acidic residues in the β-subdomain in MscS homologs from both Firmicutes and Actinobacteria (**Fig. 7a, b**). By contrast, corresponding arginine residues do not exist in the TM2 helices of MscS homologs from Gram-negative bacteria and the acidic residues in the β-subdomain are not conserved. A hydropathy plot illustrates that the conserved arginine residues of TM2 in MscS homologs from Gram-postive bacteria are located in the hydrophilic cytoplasmic extension (**Fig. 7b**, bottom panel). These findings suggest that electrostatic coupling between the cytoplasmic extension of TM2 and the β subdomain is a conserved feature of MscS homologs from Gram-positive bacterial lineages but is absent in MscS homologs from canonical Gram-negative bacteria.

**Fig. 7.**
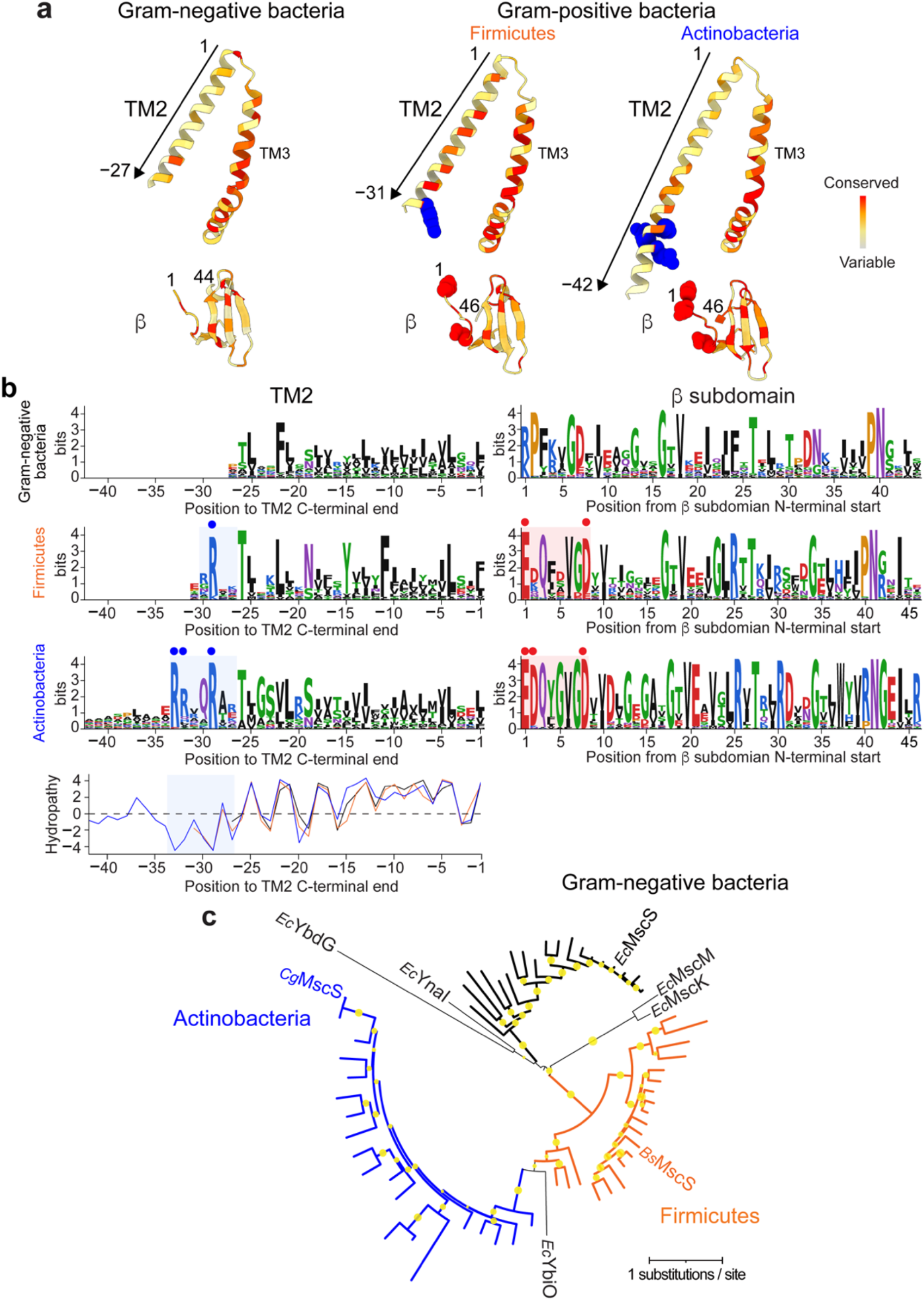
Conservation and phylogenetic relationships of MscS homologs from Gram-negative and Gram-positive bacteria. **a** Sequence conservation mapped onto the structures of representative homologs from each bacterial phylum: *Ec*MscS for Gram-negative bacteria, *Bs*MscS for Firmicutes, and *Cg*MscS for Actinobacteria. Conservation is shown as a gradient from highly variable (light gray) to highly conserved (red). Conserved arginine residues in TM2 are shown as blue spheres, and conserved acidic residues in the β subdomain are shown as red spheres. **b** Sequence-logo analyses of TM2 (left panels) and the β subdomain (right panels) of MscS homologs from Gram-negative bacteria, Firmicutes, and Actinobacteria. Logos were generated from alignments of 2,758 homologs from Gram-negative bacteria, 1,101 homologs from Firmicutes, and 817 homologs from Actinobacteria. Blue circles indicate conserved arginine residues in the cytoplasmic extension of TM2, and red circles indicate conserved acidic residues in the β subdomain. Hydropathy profiles of the consensus TM2 sequences are shown in the bottom panel. **c** Maximum-likelihood phylogenetic tree of representative MscS homologs from Gram-negative bacteria, Firmicutes, and Actinobacteria. Branches corresponding to homologs from Gram-negative bacteria, Firmicutes, and Actinobacteria are shown in black, orange, and blue, respectively. The scale bar indicates the number of amino acid substitutions per site. Circles at internal nodes indicate bootstrap support values greater than 60%.

To examine the evolutionary relationships among MscS homologs from Gram-positive and Gram-negative bacteria, we performed a phylogenetic analysis using the TM2 together with the conserved core of the MscS superfamily, comprising TM3 and the β subdomain of the cytoplasmic cage. Five MscS-like channels from *E. coli*, which contain more than three TM helices (YnaI, YbdG, YbiO, MscK, and MscM), were included as outgroups and all localized outside the three-TM-helix MscS subclades. Within the three-TM-helix channels, homologs from Firmicutes and Actinobacteria formed lineages distinct from those from Gram-negative bacteria (**Fig. 7c**). Thus, despite sharing the same overall three-TM-helix architecture, MscS homologs from Gram-positive and Gram-negative bacteria appear to have followed distinct evolutionary trajectories associated with differences in TM2–β subdomain coupling.

## DISCUSSION

MscCG was the first channel of the MscS-like superfamily to be identified in *C. glutamicum* (44) and functionally characterized (46,47,57,59), which led to the belief that it is the homolog of *Ec*MscS, a notion that did not change with the later discovery of MscCG2. However, unlike MscCG, which has four TM helices, MscCG2 has three TM helices, the same number as *Ec*MscS. Our functional analysis of MscCG2 now shows that the functional characteristics of MscCG2, the fast opening kinetics, the single-channel conductance and the desensitization/inactivation, also makes MscCG2 functionally more similar to *Ec*MscS than MscCG. MscCG has slow opening kinetics, a smaller single-channel conductance and does not desensitize/inactivate (57), characteristics that make it more similar to *E. coli* MscS-like channels, which have an MscS-like core structure but feature additional structural elements (42,65). The structural and functional characteristics together now clearly establish that MscCG2 is in fact the *Ec*MscS homolog, which is why we now refer to it as *Cg*MscS, and that MscCG is a member of the MscS-like superfamily.

A distinct difference of *Cg*MscS to *Ec*MscS is that it has a very high activation threshold, close to lytic membrane tension like *Ec*MscL (5,66,67) and *Cg*MscL (47), which raises the question why a channel with lower conductance, *Cg*MscS, is needed to open when a channel with much higher conductance, *Cg*MscL, opens at the same high membrane tension. Furthermore, *Cg*MscS also does not show gating hysteresis, meaning that it closes at a similarly high membrane tension as when it opens. Therefore, *Cg*MscS cannot play the same role as *Ec*YnaI in *E. coli*, which opens a small pore at the same high membrane tension as *Ec*MscL but then, due to is large gating hysteresis, stays open to a much lower membrane tension than *Ec*MscL (42). It thus appears that MscCG, which opens at a lower membrane tension (47), similar to that of *Ec*MscS, plays the physiological role of *Ec*MscS in *E. coli*. This view is supported by the finding that many *Corynebacteria* have lost their *Cg*MscS homologs and only express an MscCG homolog (45). It thus appears that once *Corynebacteria* acquired the MscCG channel, it made the function of *Cg*MscS redundant. It should be noted, however, that this situation is unique to *Corynebacteria*, as MscCG homologs are only expressed in *Corynebacteria*, while all other Actinobacteria express *Cg*MscS homologs. In addition, these considerations only concern the function of the MscS channels in osmoregulation and do not take into account their role in the conduction of other substrates, such as amino acids and in particular glutamate in *C. glutamicum*.

Electrophysiology revealed that *Cg*MscS generates extremely flickery currents in *E. coli* spheroplasts and SPL liposomes that prevented functional characterization of *Cg*MscS. However, the currents were substantially less flickery when electrophysiology was performed on *C. glutamicum* spheroplasts or fusion vesicles formed with native *C. glutamicum* vesicles and SPL liposomes. Since the inner membrane of *C. glutamicum* is dominated by negatively charged lipids, patch-clamp measurements were performed on SPL liposomes supplemented with negatively charged lipids, which also yielded much less flickery currents, establishing that *Cg*MscS has adapted to a highly negatively charged membrane. Negatively charged lipids could affect the open-channel stability of *Cg*MscS either through the membrane that surrounds *Cg*MscS or through the lipids in its extramembranous pockets. Our results suggest that the pocket lipids are very stably associated with *Cg*MscS and co-purify with the channel, so that they would not explain why *Cg*MscS generates more flickery currents when reconstituted into SPL liposomes with or without negatively charged lipids. It thus seems more likely that it is the physicochemical properties of the membrane surrounding *Cg*MscS that affect the flickeriness of the channel, but identifying the specific membrane characteristic(s) and the underlying mechanism(s) will require further study.

Another unresolved question concerns the structural variability of the αβ subdomain, which makes the cytoplasmic cage of *Cg*MscS distinct from those of other members of the MscS-like superfamily, which mostly show a very rigid and stable structure. The αβ subdomain of *Cg*MscS expands and contracts, independent on whether the channel is in the closed or inactivated state (**Fig. S3**, **S13** and **Movies S1**, **S2**). This structural variability is different from that recently seen in the *E. coli* MscS-like channel *Ec*MscM, in which the αβ subdomain adopts different but defined conformations based on the functional state of the channel (65). The structural variability of the αβ subdomain in *Cg*MscS is likely due to the lack of a C-terminal structural element that holds the subunits together, such as the β strand of *Ec*MscS that forms a stable β barrel at the cytoplasmic end of the cage and stabilizes the protein (68). Instead, AlphaFold3 (69) predicts, with low confidence, that the C-terminal sequence of *Cg*MscS forms an α helix, which is not predicted to form a helical bundle (**Fig. S15**). The structural variability of the αβ subdomain does not substantially affect the size of the fenestrations formed at the interface between the β and αβ subdomains but does alter the size of the axial opening of the cage. However, whether the variability in the size of the axial opening in the cytoplasmic cage is phyisiologically relevant and has an effect on single-channel conductance and/or substrate selectivity will require further study.

*Ec*MscS is the archetypal MscS channel that has been used to uncover the mechanisms underlying membrane tension-mediated channel opening of this family of MS channels, leading to the “lipids-move-first” model (23) and the understanding that channel gating in these channels involves both tilting and rotation of the TM1-2 paddles, which is then transmitted to the pore-lining TM3a helices and leads to changes in the diameter of the ion-conducting pore (14,18,19). Our structures of WT and A146V *Cg*MscS show a rotation of the TM1-2 paddles, twisting of the pore-lining TM3a helices, and a reorientation of the aromatic gate, but show no evidence that the TM1-2 paddles tilt. These observations establish a distinc gating mechanism, in which membrane tension is most likely transmitted to the pore exclusively through rotational motions within the TM domain and does not require the paddles to tilt as seen for *Ec*MscS. However, without a structure of *Cg*MscS in an open conformation, we cannot completely rule out that gating involves some tilting of the TM1-2 paddles. On the other hand, because there are no clear changes in the pocket lipids, we can rule out that the gating of *Cg*MscS is based on the lipids-move-first model.

The cytoplasmic extension of TM2 with its two conserved arginine residues that is conserved in actinobacterial MscS homologs is the structural element most likely responsible for the alternative force-transmission mechanism seen in *Cg*MscS. The two positively charged residues Arg97 and Arg101 interact with negatively charged residues Asp174 and Asp168 on the β subdomain of the cytoplasmic cage. Both the arginine residues in the TM2 extension and the aspartate residues in the β subdomain are conserved in actinobacterial MscS homologs but not in MscS homologs from Gram-negative bacteria or Firmicutes, indicating that these two salt bridges are not only conserved in actinobacterial MscS homologs but also specific to those. The requirement for these two salt bridges to break before the TM1-2 paddle can move may be the reason for the high membrane tension needed to open *Cg*MscS.

Given the high activation threshold of *Cg*MscS and the small diameter of the constriction site in the ion-conducting pathway, the structure of WT *Cg*MscS clearly represents the channel in the closed state. Assigning a functional state to the structure of A146V *Cg*MscS was less straightforward. This structure shows a rotation of the TM1-2 paddle resulting in a conformation, in which the TM1-2 paddle is now aligned with the β subdomain of the same protomer rather than with that of the neighboring protomer. Rotation of the TM1-2 paddle and the resulting aligned domain configuration is the hallmark of the open conformation of *Ec*MscS, suggesting that the structure of A146V *Cg*MscS may represent an open state. However, because the diameter of the pore is only slightly larger, 5.2 Å, than in the closed state, 4.4 Å, and because all-atom MD simulations show no meaningful ion conduction through the pore that would be consistent with an open channel, we conclude that the structure of A146V *Cg*MscS likely represents the inactivated (or potentially the desensitized) state of the channel. Notably, the structure of A146V *Cg*MscS representing the inactivated state shows the same Arg97–Asp174 and Arg101–Asp168 interactions seen in WT *Cg*MscS representing the closed state (**Fig. S16**). Even though these salt bridges now form within the same protomer rather than between adjacent protomers, the presence of the same salt bridges suggests that the closed and inactivated conformation may be similarly stable.

A morph between the closed and inactivated state of *Ec*MscS, without going through the open state, shows a substantial tilting of the TM1-2 paddles and virtually no rotation (**Movie S3**). This structural change is almost opposite from that seen in the morph between the closed (WT) and inactivated state (A146V) of *Cg*MscS, which shows almost exclusively a rotation of the TM1-2 paddles and no meaningful tilting (**Movie S4**), thus emphasizing the difference in the mechanisms underlying the gating of these two MscS homologs. Attempts to stabilize *Cg*MscS in an open conformation by βCD-mediated lipid removal from nanodiscs and delipidation by exposure to high DDM concentration did not succeed, even when the low-threshold gating A151V mutant was used. The failure of βCD treatment to induce an open conformation suggests that the mallability of the membrane-scaffold protein prevents lipid removal from creating sufficient membrane tension, a problem that is exacerbated by the likely small change in the in-plane area that occurs when *Cg*MscS transitions to the open conformation. The same issues likely prevented βCD treatment to yield a structure of YnaI in the open conformation (42). Furthermore, the flickeriness of *Cg*MscS-mediated currents and the many observed subconductance levels suggest that the open conformation of *Cg*MscS is unstable and transient, and thus difficult to stabilize under equilibrium cryo-EM conditions. This notion is supported by the fact that both the closed and inactivated conformation feature the same two salt bridges, making them thermodynamically much more stable than the in-between open conformation.

## MATERIALS AND METHODS

### Cloning and mutagenesis

The *Cg*MscS gene was amplified by PCR from the genomic DNA of industrial *Corynebacterium glutamicum* strain ATCC 13869 and cloned into a modified pEKex2-derived expression vector carrying a C-terminal 6xHis tag. The original pEKex2 vector was a gift from Prof. Reinhard Krämer (70). To optimize protein yield while maintaining cell growth, in Prof. Boris Martinac’s laboratory, the native Ptac promoter of pEKex2 was replaced with the intermediate-strength constitutive promoter I29 from a *C. glutamicum* synthetic promoter library (71), generating the modified expression plasmid pEKex2-I29. Unless otherwise stated, *Cg*MscS constructs were expressed in a mechanosensitive channel-deletion strain of *C. glutamicum*, ATCC 13032 (ΔmscCG ΔmscL), which was generated in Prof. Hisashi Kawasaki’s laboratory from the *C. glutamicum* ATCC 13032 wild-type strain (Kyowa). For expression in *Escherichia coli*, the *Cg*MscS gene was cloned into a pBAD arabinose-inducible expression vector. Single-point mutations were introduced using the Q5 Site-Directed Mutagenesis Kit (New England Biolabs) according to manufacturer’s instructions, with primers designed using NEBaseChanger. To introduce the A146V mutation, the following primers were used: *Cg*MscS-A146V-F: GGCTGGTGTTgtgCTTGGTTTTG, *Cg*MscS-A146V-R: ACACCTGCCGAGGCG. To introduce the A151V mutation, the following primers were used: *Cg*MscS-A151V-F: TGGTTTTGGCgtgCAGTCGCTGG, *Cg*MscS-A151V-R: AGCGCAACACCAGCC. All constructs were verified by DNA sequencing.

### Protein expression and purification

All *C. glutamicum* cells were cultured in Brain Heart Infusion (BHI) medium (BD BBL, 211059). Transformants generated by electroporation were selected on BHI agar plates containing 25 μg/mL kanamycin and incubated at 31.5°C. Selected colonies were used to inoculate starter cultures of 30 mL BHI medium containing 25 μg/mL kanamycin in 200 mL flasks and grown at 31.5°C. For large-scale protein expression, overnight starter cultures were used to inoculate 4 - 6 L of fresh BHI medium at a 1:200 ratio. The cultures were grown for 16 h at 18°C in the presence of 6 μg/mL kanamycin. Cells were harvested by centrifugation at 5,000 × *g* for 30 min at 4°C, and cell pellets were resuspended in cold phosphate-buffered saline (PBS), containing 10 mM disodium hydrogen phosphate, pH 7.3 to 7.5, 150 mM sodium chloride and supplemented with 50 mM betaine. The cell suspension was treated with 1 mg/mL lysozyme on ice for 5 min, followed by addition of 0.1 mg/mL DNase I, and the cells were then disrupted by sonication. Cell debris was removed by centrifugation at 16,000 × *g* for 10 min at 4°C, and membranes were collected by ultracentrifugation at 215,000 × *g* for 60 min in a Type 45 Ti fixed-angle rotor (Beckman Coulter) at 4°C. Membrane pellets were weighed and resuspended in PBS supplemented with 50 mM betaine to a concentration of 0.1 g/mL. Membrane pellets were thoroughly homogenized and 20 μL of the resulting membrane vesicles were used for liposomal fusion.

The remaining membranes were solubilized with 1% (w/v) n-dodecyl-β-D-maltopyranoside (DDM; Anatrace) for 1 - 2 h at 4°C. Solubilized membranes were clarified by centrifugation at 5,000 × *g* for 10 min at 4°C, and the supernatant was incubated with Ni-NTA affinity resin (Qiagen) for 2 h at 4°C in the presence of 5 mM imidazole. The resin was washed with 40 column volumes (CV) of PBS containing 50 mM betaine, 50 mM imidazole and 0.05% DDM, and bound protein was eluted with 10 CV of the same buffer but with 250 mM imidazole. The eluate was concentrated using an Amicon Ultra concentrator with a 100-kDa molecular-weight cut-off by centrifugation at 4,000 × *g* and 4°C. The protein was further purified by size-exclusion chromatography using a Superose 6 Increase 10/300 GL column (Cytiva) equilibrated in PBS containing 0.05% DDM. Peak fractions containing *Cg*MscS were pooled and concentrated using a 100-kDa centrifugal concentrator for cryo-EM grid preparation, SDS-PAGE analysis and liposome reconstitution, as needed. Purified protein was used immediately for cryo-EM grid freezing or stored at −80°C for later use.

### Reconstitution of *Cg*MscS into nanodiscs

The lipids 1,2-dioleoyl-sn-glycero-3-phosphocholine (DOPC), 1,2-dioleoyl-sn-glycero-3-phosphoglycerol (DOPG) and 1’,3’-bis[1,2-dioleoyl-sn-glycero-3-phospho]-glycerol (cardiolipin) were purchased from Avanti Polar Lipids and used to prepare 10 mg/mL stock solutions in PBS containing 5% DDM. For nanodisc reconstitution, purified *Cg*MscS, membrane scaffold protein MSP1E3D1 and detergent-solubilized lipids were mixed at a molar ratio of 1:20:200 and incubated for 30 min on ice. Bio-Beads SM-2 resin (Bio-Rad) was then added to 30% (v/v), and the mixture was incubated for 16 h at 4°C with gentle rotation. Bio-Beads were allowed to settle by gravity, and the supernatant was loaded onto a Superose 6 Increase 10/300 GL column equilibrated with PBS to remove empty nanodiscs and aggregated protein. Peak fractions containing *Cg*MscS-containing nanodiscs were pooled and used for cryo-EM grid preparation.

For βCD-mediated lipid removal, 0.5 mL of *Cg*MscS reconstituted into DOPG/MSP1E3D1 nanodiscs at a concentration of 0.6 mg/mL was mixed with 0.5 mL of 200 mM βCD and incubated for 16 h at 4°C. βCD-treated nanodisc samples were loaded onto a Superose 6 Increase 10/300 GL column equilibrated with PBS. Peak fractions containing *Cg*MscS-reconstituted nanodiscs were pooled and concentrated to 0.1 - 0.2 mg/mL for cryo-EM grid preparation.

### Cryo-EM grid preparation and data collection

The homogeneity of purified *Cg*MscS samples was first examined by negative-stain electron microscopy using 0.7% (w/v) uranyl formate, as described previously (72). Protein concentration was measured using a NanoDrop spectrophotometer (Thermo Fisher Scientific) and adjusted to 0.1 - 0.2 mg/mL for cryo-EM grid preparation. Aliquots of 3 μL were applied to glow-discharged Quantifoil R1.2/1.3 Cu 400 mesh or Quantifoil R2/4 Cu 300 mesh grids with graphene oxide support (Electron Microscopy Sciences). The grids were frozen with a Vitrobot Mark IV (Thermo Fisher Scientific), with the chamber maintained at 4°C and 100% humidity. After sample application, grids were blotted for 2 - 4 s with a blot force of 0 or −2 and plunge-frozen in liquid ethane cooled by liquid nitrogen.

A151V *Cg*MscS in DOPG + βCD was imaged at the New York Structural Biology Center using a 300 kV Titan Krios electron microscope (Thermos Fisher Scientific) operated with Leginon (73) and equipped with a K3 direct detector camera. Data were recorded in super-resolution counting mode at a nominal magnification of 81,000x, corresponding to a calibrated pixel size of 0.826 Å at the specimen level, and using a defocus range of −0.8 to −2.5 µm. Movies were collected at a dose rate of 20 e^−^/s/pixel. Exposures of 2 s were dose-fractionated into 40 frames of 0.05 s, resulting in 1.35 e^−^/Å^2^/frame and a total dose of 58.61 e^−^/Å^2^.

All other samples were imaged at the Cryo-EM Resource Center at the Rockefeller University using a 300-kV Titan Krios electron microscope operated with SerialEM (74) and equipped with a K3 direct detector camera and a Gatan GIF filter. All data, except for A146V *Cg*MscS in DOPG, were recorded in super-resolution counting mode at a nominal magnification of 81,000x, corresponding to a calibrated pixel size of 0.86 Å at the specimen level, and using a defocus range of −0.8 to −2.5 µm. Movies were collected at a dose rate of 20 e^−^/s/pixel. Exposures of 2 s were dose-fractionated into 40 frames of 0.05 s, resulting in 1.35 e^−^/Å^2^/frame and a total dose of 54.08 e^−^/Å^2^. Data for A146V *Cg*MscS in DOPG were recorded in counting mode at a nominal magnification of 81,000x, corresponding to a calibrated pixel size of 0.86 Å at the specimen level, and using a defocus range of −0.8 to −2.5 µm. Movies were collected at a dose rate of 30 e^−^/s/pixel. Exposures of 1.332 s were dose-fractionated into 40 frames of 0.0335 s, resulting in 1.36 e^−^/Å^2^/frame and a total dose of 54.03 e^−^/Å^2^.

Data collection parameters for all datasets are summarized in **Table S1**.

### Image processing

All image processing was performed in cryoSPARC, and unless stated otherwise, C7 symmetry was imposed in all 3D-processing steps. All movies were motion-corrected using patch motion correction, and contrast transfer function (CTF) parameters were estimated using patch CTF estimation. 2D classification was used to identify classes with averages showing structural detail, which were combined for further processing, and to discard classes with poor averages. Decoy maps, used in 3D classifications to remove junk particles, were generated using particles excluded during 2D classification and terminating 3D reconstruction jobs at an early stage. The overall resolution of maps was estimated based on the gold-standard FSC curve and the FSC = 0.143 cut-off criterion

### Cryo-EM data processing of WT *Cg*MscS in DOPG nanodiscs

A total of 8,260 movies of WT *Cg*MscS in DOPG nanodiscs were collected. After curation based on CTF values, 7,484 motion-corrected micrographs were retained for further processing. Particles were initially picked from the curated micrographs using blob picker, yielding 4,887,121 particles. These particles were extracted into 320×320-pixel boxes, binned 2-fold, and subjected to iterative 2D classification into 50 classes, which yielded 318,106 particles that were used to train an initial Topaz model. The Topaz model was used to pick particles from all curated micrographs, yielding 3,570,812 particles. These particles were extracted into 320×320-pixel boxes, binned 2-fold, and subjected to iterative 2D classification into 50 classes, which yielded 176,100 particles that were used to train a second Topaz model. This Topaz model was used to pick particles from all curated micrographs, yielding 4,007,507 particles. These particles were extracted into 320×320-pixel boxes, binned 2-fold, and subjected to iterative 2D classification into 50 classes, which yielded 327,536 particles that were combined with the final 318,106 particles from blob picking. After removing duplicate particles, the remaining particles were subjected to iterative 2D classification into 50 classes, which yielded 453,777 particles that were used to generate an *ab-initio* reconstruction.

The particles were re-extracted into 320×320-pixel boxes without binning and subjected to two rounds of heterogeneous refinement into six classes, which were seeded with three copies of the *Cg*MscS *ab-initio* map and three decoy maps. One of the *Cg*MscS target classes showed well-resolved density for both the TM and cytoplasmic domains. The 149,252 particles assigned to this class were subjected to non-uniform refinement, yielding the final map at an overall resolution of 2.9 Å.

Separately, particles assigned to all three *Cg*MscS target classes from the second heterogeneous refinement were combined, yielding 313,925 particles, and used for 3D classification and 3D variability analysis. The particles were first subjected to non-uniform refinement, followed by local refinement using a mask including the TM domain. The particles were then subjected to 3D classification without alignment into 10 classes, using a mask including the αβ subdomain and the C-terminal region. The 8 classes showing clear features for the αβ subdomain were selected. These particles were subjected to 3D variability analysis using a mask including the αβ subdomain and C-terminal region. Movies corresponding to each variability mode were generated using the simple mode in 3D Variability Display and visually evaluated.

Separately, three classes from the 3D classification representing *Cg*MscS with the most contracted, an intermediate, and the most expanded αβ subdomain were subjected to non-uniform refinement.

### Cryo-EM data processing of WT *Cg*MscS in DOPC nanodiscs

A total of 4,392 movies of WT *Cg*MscS in DOPC nanodiscs were collected. After curation based on CTF values, 3,959 motion-corrected micrographs were retained for further processing. Particles were initially picked from the curated micrographs using blob picker, yielding 2,588,361 particles. These particles were extracted into 320×320-pixel boxes, binned 2-fold, and subjected to iterative 2D classification into 50 classes, which yielded 662,720 particles that were used to train a Topaz model. The Topaz model was used to pick particles from all curated micrographs, yielding 2,036,685 particles. These particles were extracted into 320×320-pixel boxes, binned 2-fold, and subjected to iterative 2D classification into 50 classes, which yielded 753,298 particles that were combined with the final 662,720 particles from blob picking. After removing duplicate particles, the remaining particles were subjected to 2D classification into 50 classes, which yielded 778,345 particles that were used to generate an *ab-initio* reconstruction.

The particles were re-extracted into 320×320-pixel boxes without binning and subjected to a heterogeneous refinement into six classes, which was seeded with three copies of the *Cg*MscS *ab-initio* map and three decoy maps. One of the *Cg*MscS target classes showed well-resolved density for both the TM and cytoplasmic domains. The 263,723 particles assigned to this class were subjected to non-uniform refinement, yielding the final map at an overall resolution of 2.7 Å.

### Cryo-EM data processing of WT *Cg*MscS in 0.05% DDM

A total of 8,910 movies of WT *Cg*MscS solubilized in 0.05% DDM were collected. After curation based on CTF values, 8,462 motion-corrected micrographs were retained for further processing. Particles were initially picked from the curated micrographs using blob picker, yielding 5,427,356 particles. These particles were extracted into 320×320-pixel boxes, binned 2-fold, and subjected to iterative 2D classification into 50 classes, which yielded 1,248,032 particles that were used to train a Topaz model. The Topaz model was used to pick particles from all curated micrographs, yielding 4,076,940 particles. These particles were extracted into 320×320-pixel boxes, binned 2-fold, and subjected to iterative 2D classification into 50 classes, which yielded 1,470,589 particles that were combined with the final 1,248,032 particles from blob picking. After removing duplicate particles, the remaining particles were subjected to 2D classification into 50 classes, which yielded 1,197,243 particles that were used to generate an *ab-initio* reconstruction.

The particles were re-extracted into 320×320-pixel boxes without binning and subjected to a heterogeneous refinement into six classes, which was seeded with three copies of the *Cg*MscS *ab-initio* map and three decoy maps. One of the *Cg*MscS target classes showed well-resolved density for both the TM and cytoplasmic domains. The 411,926 particles assigned to this class were subjected to non-uniform refinement, yielding the final map at an overall resolution of 2.9 Å.

### Cryo-EM data processing of WT *Cg*MscS incubated in 0.5% DDM

A total of 4,664 movies of WT *Cg*MscS incubated in 0.5% DDM were collected. After curation based on CTF values, 3,837 motion-corrected micrographs were retained for further processing. Particles were initially picked from the curated micrographs using blob picker, yielding 2,828,187 particles. These particles were extracted into 320×320-pixel boxes, binned 2-fold, and subjected to iterative 2D classification into 50 classes, which yielded 314,763 particles that were used to train a Topaz model. The Topaz model was used to pick particles from all curated micrographs, yielding 2,303,751 particles. These particles were extracted into 320×320-pixel boxes, binned 2-fold, and subjected to iterative 2D classification into 50 classes, which yielded 362,050 particles that were combined with the final 314,763 particles from blob picking. After removing duplicate particles, the remaining particles were subjected to 2D classification into 50 classes, which yielded 464,062 particles that were used to generate an *ab-initio* reconstruction.

The particles were re-extracted into 320×320-pixel boxes without binning and subjected to two rounds of heterogeneous refinement into six classes, which were seeded with three copies of the *Cg*MscS *ab-initio* map and three decoy maps. One of the *Cg*MscS target classes showed well-resolved density for both the TM and cytoplasmic domains. The 147,644 particles assigned to this class were subjected to non-uniform refinement, yielding the final map an overall resolution of 2.9 Å.

### Cryo-EM data processing of A151V *Cg*MscS in DOPG nanodiscs

A total of 10,199 movies of A151V *Cg*MscS in DOPG nanodiscs were collected. After curation based on CTF values, 9,715 motion-corrected micrographs were retained for further processing. Particles were initially picked from 340 of the curated micrographs using blob picker, yielding 328,382 particles. These particles were extracted into 320×320-pixel boxes, binned 4-fold, and subjected to 2D classification into 50 classes, which yielded 214,773 particles. These particles were extracted into 320×320-pixel boxes, binned 4-fold, and subjected to 2D classification into 50 classes, which yielded 9,182 particles that were used to train a Topaz model using the same 340 curated micrographs. The Topaz model was then used to pick particles from all curated micrographs, yielding 1,975,085 particles. These particles were extracted into 320×320-pixel boxes, binned 4-fold, and subjected to iterative 2D classification into 50 classes, which yielded 597,578 particles that were used to generate an *ab-initio* reconstruction.

The particles were re-extracted into 320×320-pixel boxes without binning and subjected to heterogeneous refinement into six classes, which were seeded with three copies of the *Cg*MscS *ab-initio* map and three decoy maps. One of the *Cg*MscS target classes showed well-resolved density for both the TM and cytoplasmic domains. The 139,005 particles assigned to this class were subjected to non-uniform refinement, yielding the final map at an overall resolution of 3.0 Å.

### Cryo-EM data processing of A151V *Cg*MscS in 0.05% DDM

A total of 11,322 movies of A151V *Cg*MscS in 0.05% DDM were collected. After curation based on CTF values, 6,923 motion-corrected micrographs were retained for further processing. Particles were initially picked from the curated micrographs using blob picker, yielding 2,803,643 particles. These particles were extracted into 320×320-pixel boxes, binned 2-fold, and subjected to iterative 2D classification into 50 classes, which yielded 213,650 particles that were used to train a Topaz model. The Topaz model was then used to pick particles from all curated micrographs, yielding 3,416,739 particles. These particles were extracted into 320×320-pixel boxes, binned 2-fold, and subjected to 2D classification into 50 classes, which yielded 285,086 particles that were used to generate an *ab-initio* reconstruction.

The particles were re-extracted into 320×320-pixel boxes without binning and subjected to a heterogeneous refinement into six classes, which was seeded with three copies of the *Cg*MscS *ab-initio* map and three decoy maps. One of the *Cg*MscS target classes showed well-resolved density for both the TM and cytoplasmic domains. The 86,811 particles assigned to this class were subjected to non-uniform refinement, yielding the final map at an overall resolution of 3.4 Å.

### Cryo-EM data processing of WT *Cg*MscS in DOPG nanodiscs after βCD treatment

A total of 4,920 movies of WT *Cg*MscS in DOPG nanodiscs after βCD treatment were collected. After curation based on CTF values, 4,496 motion-corrected micrographs were retained for further processing. Particles were initially picked from 423 of the curated micrographs using blob picker, yielding 261,794 particles. These particles were extracted into 320×320-pixel boxes, binned 4-fold, and subjected to 2D classification into 50 classes, which yielded 21,202 particles that were used to train a Topaz model using the same 423 curated micrographs. The Topaz model was used to pick particles from all curated micrographs, yielding 1,430,115 particles. These particles were extracted into 320×320-pixel boxes, binned 4-fold, and subjected to iterative 2D classification into 50 classes, which yielded 345,567 particles that were used to generate an *ab-initio* reconstruction.

The particles were re-extracted into 320×320-pixel boxes without binning and subjected to a heterogeneous refinement into six classes, which was seeded with three copies of the *Cg*MscS *ab-initio* map and three decoy maps. One of the *Cg*MscS target classes showed well-resolved density for both the TM and cytoplasmic domains. The 155,171 particles assigned to this class were subjected to non-uniform refinement, yielding the final map at an overall resolution of 2.9 Å.

### Cryo-EM data processing of A151V *Cg*MscS in DOPG nanodiscs after βCD treatment

A total of 7,775 movies of A151V *Cg*MscS in DOPG nanodiscs after βCD treatment were collected. After curation based on CTF values, 7,066 motion-corrected micrographs were retained for further processing. Particles were initially picked from the curated micrographs using blob picker, yielding 4,828,825 particles. These particles were extracted into 320×320-pixel boxes, binned 2-fold, and subjected to iterative 2D classification into 50 classes, which yielded 467,151 particles that were used to train a Topaz model. The Topaz model was used to pick particles from all curated micrographs, yielding 3,656,769 particles. These particles were extracted into 320×320-pixel boxes, binned 2-fold, and subjected to iterative 2D classification into 50 classes, which yielded 582,258 particles that were combined with the final 467,151 particles from blob picking. After removing duplicate particles, the remaining particles were subjected to 2D classification into 50 classes, which yielded 696,903 particles that were used to generate an *ab-initio* reconstruction.

The particles were re-extracted into 320×320-pixel boxes without binning and subjected to heterogeneous refinement into six classes, which was seeded with three copies of the *Cg*MscS *ab-initio* map and three decoy maps. One of the *Cg*MscS target classes showed well-resolved density for both the TM and cytoplasmic domains. The 302,339 particles assigned to this class were subjected to non-uniform refinement, yielding the final map at an overall resolution of 2.8 Å.

### Cryo-EM data processing of A146V *Cg*MscS in DOPG nanodiscs

A total of 11,457 movies of A146V *Cg*MscS in DOPG nanodiscs were collected. After curation based on CTF values, 8,325 motion-corrected micrographs were retained for further processing. Particles were initially picked from the curated micrographs using blob picker, yielding 4,448,182 particles. These particles were extracted into 320×320-pixel boxes, binned 2-fold, and subjected to iterative 2D classification into 50 classes, which yielded 496,450 particles that were used to train a Topaz model. The Topaz model was used to pick particles from all curated micrographs, yielding 4,433,364 particles. These particles were extracted into 320×320-pixel boxes, binned 2-fold, and subjected to iterative 2D classification into 50 classes, which yielded 979,754 particles that were combined with the final 496,450 particles from blob picking. After removing duplicate particles, the remaining particles were subjected to 2D classification into 50 classes, which yielded 1,104,819 particles that were used to generate an *ab-initio* reconstruction.

The particles were re-extracted into 320×320-pixel boxes without binning and subjected to heterogeneous refinement into six classes, which was seeded with three copies of the *Cg*MscS *ab-initio* target map and three decoy maps. One of the *Cg*MscS target classes showed well-resolved density for both the TM and cytoplasmic domains. The 266,481 particles assigned to this class were subjected to non-uniform refinement, yielding the final map at an overall resolution of 3.3 Å.

Separately, particles assigned to all three *Cg*MscS target classes from the second heterogeneous refinement were combined, yielding 665,814 particles, and used for 3D classification and 3D variability analysis. The particles were first subjected to non-uniform refinement, followed by local refinement using a mask including the TM domain. The particles were then subjected to 3D classification without alignment into 10 classes, using a mask including the αβ subdomain and the C-terminal region. The 5 classes showing clear features for the αβ subdomain were selected. These particles were subjected to 3D variability analysis using a mask including the αβ subdomain and C-terminal region. Movies corresponding to each variability mode were generated using the simple mode in 3D Variability Display and visually evaluated.

Separately, three classes from the 3D classification representing A146V *Cg*MscS with the most contracted, an intermediate, and the most expanded αβ subdomain were selected to non-uniform refinement.

### Model building and refinement

For model building of WT *Cg*MscS in DDM, a model of *Cg*MscS generated with AlphaFold3 (69) was rigid-body docked into the cryo-EM density map using UCSF ChimeraX (75). One monomer was manually adjusted and refined in Coot (76). Densities corresponding to pocket lipids were initially modeled using dioleoyl phosphatidylglycerol. Terminal atoms of the acyl chains and headgroups poorly supported by the density were removed. The protein/lipid model for one monomer was further refined using phenix.real_space_refine (77). Seven copies of the refined monomer were then docked into the cryo-EM map, followed by real-space refinement of the complete heptameric model to remove steric clashes and to optimize intersubunit contacts. The WT *Cg*MscS DDM model was used as starting point to build models into all other density maps. For each map, the model was first rigid-body docked into the map in UCSF ChimeraX, followed by iterative rounds of manual adjustment in Coot and refinement using phenix.real_space_refine. Cryo-EM maps were sharpened using DeepEMhancer (78) when needed to improve map interpretation. Regions with weak or discontinuous density, including poorly resolved loops and terminal residues, were omitted from the final models. All final models were validated using phenix.validation_cryoem. Refinement and validation statistics are summarized in **Table S1**.

Pore radii were calculated using HOLE v2.2 (62), and pore-radius profiles were determined along the channel axis by calculating the distance between the centre of the pore and the pore surface at each point along the axis.

### Preparation of *E. coli* giant spheroplasts

WT *Cg*MscS was expressed using the pBAD plasmid in *E. coli* MJF612(DE3) cells (Frag1, Δ*mscL*::cm, Δ*kefA*::kan, Δ*ybdG*::apr, Δ*yggB*) (58). Giant spheroplasts were prepared as described previously (3), with slight modifications. Briefly, 100 μL of an overnight culture was transferred to 10 mL LB medium containing 50 μg/mL ampicillin (Research Products International) and incubated at 37°C until OD₆₀₀ reached 0.5 - 1.0. To induce bacterial filament formation, 3 mL of this culture was transferred to 27 mL LB medium containing 50 μg/mL ampicillin and 180 μL of 10 mg/mL cephalexin (Sigma-Aldrich), corresponding to a final cephalexin concentration of 60 μg/mL. The culture was incubated at 37 °C for 60 - 90 min and when filaments reached approximately 100 μm in length, as assessed by light microscopy, protein expression was induced by adding arabinose to a final concentration of 0.05% (w/v). After incubation for 1 h at 37°C, cells were harvested by centrifugation at 3,000 × *g* for 10 min at 4°C and resuspended in 2.5 mL of 0.8 M sucrose. To generate giant spheroplasts, the following solutions were added sequentially: 1) 150 μL of 1 M Tris-HCl, pH 8.0; 2) 120 μL of 5 mg/mL lysozyme (OmniPur, Calbiochem); 3) 50 μL of 5 mg/mL DNase I (Worthington); and 4) 150 μL of 125 mM EDTA, pH 8.0, adjusted with NaOH. The final concentrations were approximately 50 mM Tris-HCl, 0.2 mg/mL lysozyme, 0.084 mg/mL DNase I, and 6.3 mM EDTA. After incubation for 10 - 13 min at room temperature, the reaction was stopped by adding stop solution containing 0.8 M sucrose, 10 mM Tris-HCl, pH 8.0, and 20 mM MgCl₂. To separate giant spheroplasts from cell debris, 2 mL of the cell suspension was carefully layered onto 7 mL of cushion solution containing 0.8 M sucrose, 10 mM Tris-HCl, pH 8.0, and 10 mM MgCl₂ in a 15-mL Falcon tube. Samples were centrifuged at 300 × *g* for 5 min at 4°C. The supernatant was carefully removed by aspiration, leaving 1 mL to resuspend the pellet. Aliquots of giant spheroplast suspension were stored at −20°C until use.

### Preparation of *Cg*MscS proteoliposomes and fusion vesicles

Soy polar lipids (SPL; Avanti) were dissolved in chloroform and dried under a gentle stream of nitrogen gas. The resulting lipid film was resuspended at 10 mg/mL in dehydration–rehydration buffer containing 5 mM HEPES, pH 7.2, adjusted with KOH, and 200 mM KCl. The lipid suspension was vortexed and sonicated in a water bath for 15 min at room temperature.

For proteoliposome preparation, purified *Cg*MscS was added to the SPL suspension at a protein-to-lipid ratio of 1:50 (w/w), and the mixture was incubated for 1 h at room temperature with agitation. The sample volume was increased to 3 mL with dehydration–rehydration buffer to reduce the detergent concentration, and detergent was removed by incubation with 100 mg Bio-Beads SM-2 for 3 h at room temperature. Bio-Beads were allowed to settle by gravity, and the supernatant was transferred to an ultracentrifuge tube. Proteoliposomes were pelleted by ultracentrifugation with a TLA-100.3 rotor (Beckman Coulter) at 100,000 × *g* for 30 min at 4°C. The protein/lipid pellet was resuspended, spotted onto a glass microscope slide, and desiccated overnight at room temperature. The resulting dried film was rehydrated overnight in dehydration– rehydration buffer at room temperature before being used in patch-clamp recordings.

For the preparation of fusion vesicles containing *Ec*MscS, *Cg*MscS, or MscCG, 2 mg of *Ec*MscS, *Cg*MscS, or MscCG-overexpressing *C. glutamicum* membrane vesicles prepared as described previously (59) were mixed with 200 μL of 10 mg/mL SPL liposomes in dehydration– rehydration buffer. The mixture was incubated for 1 h at room temperature, spotted onto a glass microscope slide, and vacuum-desiccated overnight. The resulting dried film was rehydrated with dehydration–rehydration buffer for 3 h to overnight at 4°C before use in patch-clamp recordings.

### Patch-clamp electrophysiology

Borosilicate glass pipettes (Drummond) were pulled using a Sutter P-97 Flaming/Brown micropipette puller. The pipette tip size was adjusted by changing the velocity parameter of the puller to obtain a resistance of 1.5 - 2.5 MΩ in the recording solution. Pipette resistance was measured using the membrane test function of an Axopatch 200B amplifier (Molecular Devices) operated with Clampex 11.2 software. All recordings were performed in the inside-out excised patch configuration.

For recordings from proteoliposomes and fusion vesicles, symmetrical bath and pipette solutions were used, containing 5 mM HEPES, pH 7.2, adjusted with KOH, 200 mM KCl, and 40 mM MgCl_2_. Single-channel currents were amplified using an Axopatch 200B amplifier, sampled at 5 kHz and low-pass filtered at 1 kHz using a Digidata 1550B digitizer. Negative pressure was applied to the recording pipette using a High Speed Pressure Clamp-1 system (ALA Scientific Instruments) and monitored with a pressure gauge (World Precision Instruments). Pressure was applied either manually using a syringe or automatically using a pressure/vacuum pump unit connected to the high-speed pressure-clamp system.

For recordings from *E. coli* giant spheroplasts, the pipette solution contained 5 mM HEPES, pH 7.2, adjusted with KOH, 200 mM KCl, and 90 mM MgCl_2_, and 2 mM CaCl_2_. The bath solution contained the same components supplemented with 400 mM sucrose to prevent osmotic changes during recording. A 10-μL aliquot of giant spheroplast suspension was placed onto the surface of the bath solution and allowed to settle to the bottom of the recording chamber. Cell debris was removed from the bath surface by gentle pipetting before recordings.

### All-atom molecular-dynamics (MD) simulations

Systems for all-atom MD simulations of the structures of *Ec*MscS in the closed (PDB: 6VYK) and open (PDB: 2VV5) conformations and the structures of WT *Cg*MscS and the A146V mutant were built using the CHARMM-GUI Membrane Builder web server (79,80). To minimize the size of the simulation systems, the αβ subdomains of *Ec*MscS (a.a. 181-286) and *Cg*MscS (a.a. 220-334) were not included in the simulations. The *Ec*MscS structures were placed in a membrane composed of 1-palmitoyl-2-oleoyl-sn-glycero-3-phosphoethanolamine (POPE), while the *Cg*MscS structures were placed in a membrane composed of 1-palmitoyl-2-oleoyl-sn-glycero-3-phosphoglycerol (POPG). In the system of *Ec*MscS in the closed conformation, two POPE lipids were manually placed in the vicinity of the pore of the channel to serve as pore lipids. Each structure was solvated in TIP3P water and 500 mM KCl (81,82). The van der Waals interactions were smoothly switched off between 10 and 12 Å using the force-switch method (83). The Particle Mesh Ewald method was used to model the long-range electrostatic interactions with a cut-off value of 1.2 nm, a Fourier grid spacing of 0.12 nm, and an interpolation order of 4 for the Ewald mesh (84,85). Temperature and pressure were maintained at 310.15 K and 1 atm using the v-rescale thermostat and C-rescale barostat, respectively (86,87). Simulations were run with a 2-fs time step. The LINCS algorithm was used to constrain hydrogen bonds (88). All simulations were run with the GROMACS 2025 software package and the CHARMM36m force field (89–91).

The standard 6-step CHARMM-GUI equilibration protocol was used to equilibrate the system. The final equilibration step that restrains only the protein backbone with a force constant of 50 kJ mol^−1^ nm^−2^ was extended to a total of 100 ns to allow lipids to associate with the hydrophobic pockets of the channels. To investigate ion conduction through the pores of *Ec*MscS and *Cg*MscS, simulations were performed using a constant electric field of approximately 200 mV for 200 ns while restraining the protein backbone with a 50 kJ mol^−1^ nm^−2^ force constant (92). All simulations were replicated five times.

Ion permeation through the pore of the channels was analyzed using MDAnalysis (93,94). A cylindrical region that represented the dimensions of the pore of each channel formed by the TM3a helices was defined. Only ions that passed through one side of that region to the other were counted as ion-permeation events.

### Phylogenetic analysis and conservation mapping

Bacterial 3TM-type MscS homologs were identified by BLASTP searches using *Ec*MscS, *Bacillus subtilis* MscS (*Bs*MscS) and *Cg*MscS as representative query sequences for MscS homologs from Gram-negative bacteria, Firmicutes and Actinobacteria, respectively. Retrieved sequences were filtered to remove fragments, duplicate entries and proteins with incomplete MscS-superfamily core regions. TM topology was predicted with DeepTMHMM, and proteins predicted to contain the canonical three-TM-helix architecture were retained for subsequent analyses. Sequences were classified according to taxonomic lineage, yielding final datasets of 6,505 homologs for Gram-negative bacteria, 2,031 homologs for Firmicutes, and 1,414 homologs for Actinobacteria.

For the phylogenetic analysis, representative sequences (2,758 for Gram-negative bacteria, 1,101 for Firmicutes, and 817 for Actinobacteria) were selected from each lineage to reduce redundancy and avoid over-representation of closely related genera. For these homologs, the regions spanning TM2, TM3 and the β subdomain of the cytoplasmic cage were extracted, subjected to multiple-sequence alignments with MAFFT (95), and poorly aligned positions were removed with trimAl (96). Maximum-likelihood trees were inferred with IQ-TREE (97) using automatic substitution model selection and branch support estimation. The resulting tree was rooted using the five MscS-like channels from *E. coli* that contain more than three TM helices (YnaI, YbdG, YbiO, MscK and MscM). Trees were visualized and annotated with iTOL (98), with clade assignments, TM2-extension status and average protein length displayed as annotation layers.

For conservation mapping, lineage-specific alignments were generated separately for homologs from Gram-negative bacteria, Firmicutes and Actinobacteria. Conservation scores were calculated for each alignment position using Jalview (99) and mapped onto representative structures for each lineage, namely a cryo-EM structure of *Ec*MscS for Gram-negative bacteria (PDB: 6VYK) (22), an AlphaFold prediction of *Bs*YkuT for Firmicutes, and the cryo-EM structure of *Cg*MscS in DOPG nanodiscs for Actinobacteria. Conservation values were displayed as a color gradient from variable to conserved residues. This analysis was used to compare conservation patterns within TM2 and to identify residues enriched within lineage-specific structural features.

For the logo analysis (100) of the TM2 sequence, TM2 boundaries were defined using the structural models of *Ec*MscS, *Bs*MscS and *Cg*MscS. Confidently aligned TM2–TM3a regions were extracted and realigned separately for each lineage. For the logo analysis of the β subdomain sequence, TM2–TM3 and the β subdomain were defined using the structural models of *Ec*MscS, *Bs*MscS and *Cg*MscS. Confidently aligned regions were extracted and realigned separately for each lineage. The final logo datasets contained 2,758 MscS sequences from Gram-negative bacteria, 1,101 sequences from Firmicutes, and 817 sequences from Actinobacteria. These alignments were then used to generate clade-specific sequence logos. The resulting consensus sequence for TM2 of each group was analyzed for hydropathy profiles. Conserved arginine residues in the TM2 of Firmicutes and Actinobacteria homologs were mapped onto the *Cg*MscS structure and compared with the positions of aspartate residues in the β subdomain of the cytoplasmic cage to assess potential electrostatic coupling between the sensor paddle and cytoplasmic cage.

## Data availability

The cryo-EM maps have been deposited in the Electron Microscopy Data Bank under accession codes EMD-78718 (WT *Cg*MscS in DOPG nanodiscs), EMD-78719 (WT *Cg*MscS in DOPC nanodiscs), EMD-78655 (WT *Cg*MscS in 0.05% DDM), EMD-78729 (WT *Cg*MscS in 0.5% DDM), EMD-78733 (A151V *Cg*MscS in DOPG nanodiscs), EMD-78734 (A151V *Cg*MscS in 0.05% DDM), EMD-78720 (WT *Cg*MscS in DOPG nanodiscs treated with βCD), EMD-78732 (A151V *Cg*MscS in DOPG nanodiscs treated with βCD), and EMD-78730 (A146V *Cg*MscS in DOPG nanodiscs). The atomic coordinates have been deposited in the Protein Data Bank under accession codes 38CI (WT *Cg*MscS in DOPG nanodiscs), 38CJ (WT *Cg*MscS in DOPC nanodiscs), 37ZB (WT *Cg*MscS in 0.05% DDM), 38CN (WT *Cg*MscS in 0.5% DDM), 38CW (A151V *Cg*MscS in DOPG nanodiscs), 38CX (A151V *Cg*MscS in 0.05% DDM), 38CK (WT *Cg*MscS in DOPG nanodiscs treated with βCD), 38CV (A151V *Cg*MscS in DOPG nanodiscs treated with βCD), and 38CO (A146V *Cg*MscS in DOPG nanodiscs).

## ACKNOWLEDGEMENTS

We thank M. Ebrahim, J. Sotiris, and H. Ng at the Evelyn Gruss Lipper Cryo-EM Resource Center of The Rockefeller University for assistance with cryo-EM data collection and members of the Walz group for helpful discussions. We thank Prof Hisashi Kawasaki for providing industrial *Corynebacterium glutamicum* strain ATCC 13869. Some of this work was performed at the Simons Electron Microscopy Center at the New York Structural Biology Center, with major support from the Simons Foundation (SF349247).

## Author contributions

Y.N. and T.W. conceived the study. T.W. supervised the research. Y.N. performed the cryo-EM and patch-clamp experiments and the sequence conservation and phylogenetic analyses. G.H. performed the MD simulations. Y.N. and G.H. analyzed the results. Y.N. prepared the manuscript with assistance of T.W. All authors reviewed and agreed with the manuscript.

## References

1. Kefauver JM, Ward AB, Patapoutian A. Discoveries in structure and physiology of mechanically activated ion channels. Nature. 2020 Nov 26;587(7835):567–76. doi:10.1038/s41586-020-2933-1

2. Sukharev SI, Martinac B, Arshavsky VY, Kung C. Two types of mechanosensitive channels in the Escherichia coli cell envelope: solubilization and functional reconstitution. Biophys J. 1993 Jul;65(1):177–83. doi:10.1016/S0006-3495(93)81044-0

3. Martinac B, Buechner M, Delcour AH, Adler J, Kung C. Pressure-sensitive ion channel in Escherichia coli. Proc Natl Acad Sci. 1987 Apr;84(8):2297–301. doi:10.1073/pnas.84.8.2297

4. Sukharev S. Purification of the Small Mechanosensitive Channel of Escherichia coli (MscS): the Subunit Structure, Conduction, and Gating Characteristicsin Liposomes. Biophys J. 2002 Jul;83(1):290–8. doi:10.1016/S0006-3495(02)75169-2

5. Nomura T, Cranfield CG, Deplazes E, Owen DM, Macmillan A, Battle AR, et al. Differential effects of lipids and lyso-lipids on the mechanosensitivity of the mechanosensitive channels MscL and MscS. Proc Natl Acad Sci. 2012 May 29;109(22):8770–5. doi:10.1073/pnas.1200051109

6. Sukharev SI, Blount P, Martinac B, Blattner FR, Kung C. A large-conductance mechanosensitive channel in E. coli encoded by mscL alone. Nature. 1994 Mar;368(6468):265–8. doi:10.1038/368265a0

7. Martinac B, Adler J, Kung C. Mechanosensitive ion channels of E. coli activated by amphipaths. Nature. 1990 Nov;348(6298):261–3. doi:10.1038/348261a0

8. Britt M, Moller E, Maramba J, Anishkin A, Sukharev S. MscS inactivation and recovery are slow voltage-dependent processes sensitive to interactions with lipids. Biophys J. 2024 Jan;123(2):195–209. doi:10.1016/j.bpj.2023.12.007

9. Belyy V, Kamaraju K, Akitake B, Anishkin A, Sukharev S. Adaptive behavior of bacterial mechanosensitive channels is coupled to membrane mechanics. J Gen Physiol. 2010 Jun 1;135(6):641–52. doi:10.1085/jgp.200910371

10. Cox CD, Nomura T, Ziegler CS, Campbell AK, Wann KT, Martinac B. Selectivity mechanism of the mechanosensitive channel MscS revealed by probing channel subconducting states. Nat Commun. 2013 Jul 10;4(1):2137. doi:10.1038/ncomms3137

11. Akitake B, Anishkin A, Liu N, Sukharev S. Straightening and sequential buckling of the pore-lining helices define the gating cycle of MscS. Nat Struct Mol Biol. 2007 Dec;14(12):1141–9. doi:10.1038/nsmb1341

12. Edwards MD, Bartlett W, Booth IR. Pore Mutations of the Escherichia coli MscS Channel Affect Desensitization but Not Ionic Preference. Biophys J. 2008 Apr;94(8):3003–13. doi:10.1529/biophysj.107.123448

13. Edwards MD, Li Y, Kim S, Miller S, Bartlett W, Black S, et al. Pivotal role of the glycine-rich TM3 helix in gating the MscS mechanosensitive channel. Nat Struct Mol Biol. 2005 Feb;12(2):113–9. doi:10.1038/nsmb895

14. Bass RB, Strop P, Barclay M, Rees DC. Crystal Structure of *Escherichia coli* MscS, a Voltage-Modulated and Mechanosensitive Channel. Science. 2002 Nov 22;298(5598):1582–7. doi:10.1126/science.1077945

15. Perozo E, Rees DC. Structure and mechanism in prokaryotic mechanosensitive channels. Curr Opin Struct Biol. 2003 Aug;13(4):432–42. doi:10.1016/S0959-440X(03)00106-4

16. Sotomayor M, Schulten K. Molecular Dynamics Study of Gating in the Mechanosensitive Channel of Small Conductance MscS. Biophys J. 2004 Nov;87(5):3050–65. doi:10.1529/biophysj.104.046045

17. Anishkin A, Sukharev S. Water Dynamics and Dewetting Transitions in the Small Mechanosensitive Channel MscS. Biophys J. 2004 May;86(5):2883–95. doi:10.1016/S0006-3495(04)74340-4

18. Wang W, Black SS, Edwards MD, Miller S, Morrison EL, Bartlett W, et al. The Structure of an Open Form of an *E. coli* Mechanosensitive Channel at 3.45 Å Resolution. Science. 2008 Aug 29;321(5893):1179–83. doi:10.1126/science.1159262

19. Flegler VJ, Rasmussen A, Borbil K, Boten L, Chen HA, Deinlein H, et al. Mechanosensitive channel gating by delipidation. Proc Natl Acad Sci. 2021 Aug 17;118(33):e2107095118. doi:10.1073/pnas.2107095118

20. Rasmussen T, Flegler VJ, Rasmussen A, Böttcher B. Structure of the Mechanosensitive Channel MscS Embedded in the Membrane Bilayer. J Mol Biol. 2019 Aug;431(17):3081–90. doi:10.1016/j.jmb.2019.07.006

21. Reddy B, Bavi N, Lu A, Park Y, Perozo E. Molecular basis of force-from-lipids gating in the mechanosensitive channel MscS. eLife. 2019 Dec 27;8:e50486. doi:10.7554/eLife.50486

22. Zhang Y, Daday C, Gu RX, Cox CD, Martinac B, De Groot BL, et al. Visualization of the mechanosensitive ion channel MscS under membrane tension. Nature. 2021 Feb 18;590(7846):509–14. doi:10.1038/s41586-021-03196-w

23. Pliotas C, Dahl ACE, Rasmussen T, Mahendran KR, Smith TK, Marius P, et al. The role of lipids in mechanosensation. Nat Struct Mol Biol. 2015 Dec;22(12):991–8. doi:10.1038/nsmb.3120

24. Levina N. Protection of Escherichia coli cells against extreme turgor by activation of MscS and MscL mechanosensitive channels: identification of genes required for MscS activity. EMBO J. 1999 Apr 1;18(7):1730–7. doi:10.1093/emboj/18.7.1730

25. Çetiner U, Rowe I, Schams A, Mayhew C, Rubin D, Anishkin A, et al. Tension-activated channels in the mechanism of osmotic fitness in *Pseudomonas aeruginosa*. J Gen Physiol. 2017 May 1;149(5):595–609. doi:10.1085/jgp.201611699

26. Ramsey K, Britt M, Maramba J, Ushijima B, Moller E, Anishkin A, et al. The dynamic hypoosmotic response of Vibrio cholerae relies on the mechanosensitive channel MscS. iScience. 2024 Jun;27(6):110001. doi:10.1016/j.isci.2024.110001

27. Brito LF, Luciano D, Irla M, Virant D, Courtade G, Brautaset T. Identification of MscS as a Key L-Glutamate Exporter in Bacillus methanolicus. Microb Biotechnol. 2025 Oct;18(10):e70252. doi:10.1111/1751-7915.70252 PubMed PMID: 41123049; PubMed Central PMCID: PMC12541555.

28. Kloda A, Martinac B. Mechanosensitive Channels in Archaea. Cell Biochem Biophys. 2001;34(3):349–81. doi:10.1385/CBB:34:3:349

29. Nakayama Y, Yoshimura K, Iida H. Organellar mechanosensitive channels in fission yeast regulate the hypo-osmotic shock response. Nat Commun. 2012 Aug 21;3(1):1020. doi:10.1038/ncomms2014

30. Haswell ES, Meyerowitz EM. MscS-like Proteins Control Plastid Size and Shape in Arabidopsis thaliana. Curr Biol. 2006 Jan;16(1):1–11. doi:10.1016/j.cub.2005.11.044

31. Jojoa-Cruz S, Saotome K, Tsui CCA, Lee WH, Sansom MSP, Murthy SE, et al. Structural insights into the Venus flytrap mechanosensitive ion channel Flycatcher1. Nat Commun. 2022 Feb 14;13(1):850. doi:10.1038/s41467-022-28511-5

32. Suda H, Asakawa H, Hagihara T, Ohi S, Segami S, Hasebe M, et al. MSL10 is a high-sensitivity mechanosensor in the tactile sense of the Venus flytrap. Nat Commun. 2025 Sep 30;16(1):8280. doi:10.1038/s41467-025-63419-w

33. Dave N, Cetiner U, Arroyo D, Fonbuena J, Tiwari M, Barrera P, et al. A novel mechanosensitive channel controls osmoregulation, differentiation, and infectivity in Trypanosoma cruzi. eLife. 2021 Jul 2;10:e67449. doi:10.7554/eLife.67449

34. Pivetti CD, Yen MR, Miller S, Busch W, Tseng YH, Booth IR, et al. Two Families of Mechanosensitive Channel Proteins. Microbiol Mol Biol Rev. 2003 Mar;67(1):66–85. doi:10.1128/MMBR.67.1.66-85.2003

35. Malcolm HR, Maurer JA. The Mechanosensitive Channel of Small Conductance (MscS) Superfamily: Not Just Mechanosensitive Channels Anymore. ChemBioChem. 2012 Sep 24;13(14):2037–43. doi:10.1002/cbic.201200410

36. Berg A, Berntsson RPA, Barandun J. Nematocida displodere mechanosensitive ion channel of small conductance 2 assembles into a unique 6-channel super-structure in vitro. Silman I, editor. PLOS ONE. 2024 Jul 22;19(7):e0301951. doi:10.1371/journal.pone.0301951

37. Zhang J, Bhatt A, Maksaev G, Luo YL, Yuan P. Lipid-mediated gating of a miniature mechanosensitive MscS channel from Trypanosoma cruzi. Nat Commun. 2025 Aug 8;16(1):7339. doi:10.1038/s41467-025-62757-z

38. Mount J, Maksaev G, Summers BT, Fitzpatrick JAJ, Yuan P. Structural basis for mechanotransduction in a potassium-dependent mechanosensitive ion channel. Nat Commun. 2022 Nov 12;13(1):6904. doi:10.1038/s41467-022-34737-0

39. Flegler VJ, Rasmussen T, Böttcher B. How Functional Lipids Affect the Structure and Gating of Mechanosensitive MscS-like Channels. Int J Mol Sci. 2022 Dec 1;23(23):15071. doi:10.3390/ijms232315071

40. Flegler VJ, Rasmussen A, Hedrich R, Rasmussen T, Böttcher B. Mechanosensitive channel engineering: A study on the mixing and matching of YnaI and MscS sensor paddles and pores. Nat Commun. 2025 Aug 23;16(1):7881. doi:10.1038/s41467-025-63253-0

41. Edwards MD, Black S, Rasmussen T, Rasmussen A, Stokes NR, Stephen TL, et al. Characterization of three novel mechanosensitive channel activities in *Escherichia coli*. Channels. 2012 Jul;6(4):272–81. doi:10.4161/chan.20998

42. Will N, Hiotis G, Nakayama Y, Angiulli G, Zhou Z, Cox CD, et al. Lipid interactions and gating hysteresis suggest a physiological role for mechanosensitive channel YnaI. Nat Commun. 2025 Aug 12;16(1):7472. doi:10.1038/s41467-025-62805-8

43. Cox CD, Nakayama Y, Nomura T, Martinac B. The evolutionary ‘tinkering’ of MscS-like channels: generation of structural and functional diversity. Pflüg Arch - Eur J Physiol. 2015 Jan;467(1):3–13. doi:10.1007/s00424-014-1522-2

44. Nakamura J, Hirano S, Ito H, Wachi M. Mutations of the *Corynebacterium glutamicum* NCgl1221 Gene, Encoding a Mechanosensitive Channel Homolog, Induce L -Glutamic Acid Production. Appl Environ Microbiol. 2007 Jul 15;73(14):4491–8. doi:10.1128/AEM.02446-06

45. Wang Y, Cao G, Xu D, Fan L, Wu X, Ni X, et al. A Novel Corynebacterium glutamicum L-Glutamate Exporter. Vieille C, editor. Appl Environ Microbiol. 2018 Mar 15;84(6):e02691–17. doi:10.1128/AEM.02691-17

46. Börngen K, Battle AR, Möker N, Morbach S, Marin K, Martinac B, et al. The properties and contribution of the Corynebacterium glutamicum MscS variant to fine-tuning of osmotic adaptation. Biochim Biophys Acta BBA - Biomembr. 2010 Nov;1798(11):2141–9. doi:10.1016/j.bbamem.2010.06.022

47. Nakayama Y, Komazawa K, Bavi N, Hashimoto Kichi, Kawasaki H, Martinac B. Evolutionary specialization of MscCG, an MscS-like mechanosensitive channel, in amino acid transport in Corynebacterium glutamicum. Sci Rep. 2018 Aug 27;8(1):12893. doi:10.1038/s41598-018-31219-6

48. Nakayama Y, Yoshimura K, Iida H. A Gain-of-Function Mutation in Gating of Corynebacterium glutamicum NCgl1221 Causes Constitutive Glutamate Secretion. Appl Environ Microbiol. 2012 Aug;78(15):5432–4. doi:10.1128/AEM.01310-12

49. Becker M, Börngen K, Nomura T, Battle AR, Marin K, Martinac B, et al. Glutamate efflux mediated by Corynebacterium glutamicum MscCG, Escherichia coli MscS, and their derivatives. Biochim Biophys Acta BBA - Biomembr. 2013 Apr;1828(4):1230–40. doi:10.1016/j.bbamem.2013.01.001

50. Kawasaki H, Martinac B. Mechanosensitive channels of Corynebacterium glutamicum functioning as exporters of l-glutamate and other valuable metabolites. Curr Opin Chem Biol. 2020 Dec;59:77–83. doi:10.1016/j.cbpa.2020.05.005

51. Nie Z, Liu P, Wang Y, Guo X, Tan Z, Shen J, et al. Directed Evolution and Rational Design of Mechanosensitive Channel MscCG2 for Improved Glutamate Excretion Efficiency. J Agric Food Chem. 2021 Dec 29;69(51):15660–9. doi:10.1021/acs.jafc.1c07086

52. Zheng W, Wuyun Q, Li Y, Liu Q, Zhou X, Peng C, et al. Deep-learning-based single-domain and multidomain protein structure prediction with D-I-TASSER. Nat Biotechnol. 2026 Apr;44(4):641–53. doi:10.1038/s41587-025-02654-4

53. Nie Z, Liu P, Yew M, Shen J, Sun J, Schwaneberg U, et al. Channel Engineering of a Glutamate Exporter. ChemBioChem. 2025 Jan 14;26(2):e202400540. doi:10.1002/cbic.202400540

54. Özcan N, Ejsing CS, Shevchenko A, Lipski A, Morbach S, Krämer R. Osmolality, Temperature, and Membrane Lipid Composition Modulate the Activity of Betaine Transporter BetP in *Corynebacterium glutamicum*. J Bacteriol. 2007 Oct 15;189(20):7485–96. doi:10.1128/JB.00986-07

55. Nagakubo T, Tahara YO, Miyata M, Nomura N, Toyofuku M. Mycolic acid-containing bacteria trigger distinct types of membrane vesicles through different routes. iScience. 2021 Jan;24(1):102015. doi:10.1016/j.isci.2020.102015

56. Belyy V, Anishkin A, Kamaraju K, Liu N, Sukharev S. The tension-transmitting “clutch” in the mechanosensitive channel MscS. Nat Struct Mol Biol. 2010 Apr;17(4):451–8. doi:10.1038/nsmb.1775

57. Nakayama Y, Yoshimura K, Iida H. Electrophysiological Characterization of the Mechanosensitive Channel MscCG in Corynebacterium glutamicum. Biophys J. 2013 Sep;105(6):1366–75. doi:10.1016/j.bpj.2013.06.054

58. Schumann U, Edwards MD, Rasmussen T, Bartlett W, Van West P, Booth IR. YbdG in *Escherichia coli* is a threshold-setting mechanosensitive channel with MscM activity. Proc Natl Acad Sci. 2010 Jul 13;107(28):12664–9. doi:10.1073/pnas.1001405107

59. Nakayama Y, Rohde PR, Martinac B. “Force-From-Lipids” Dependence of the MscCG Mechanosensitive Channel Gating on Anionic Membranes. Microorganisms. 2023 Jan 12;11(1):194. doi:10.3390/microorganisms11010194

60. Kamaraju K, Belyy V, Rowe I, Anishkin A, Sukharev S. The pathway and spatial scale for MscS inactivation. J Gen Physiol. 2011 Jul 1;138(1):49–57. doi:10.1085/jgp.201110606

61. Nomura T, Cox CD, Bavi N, Sokabe M, Martinac B. Unidirectional incorporation of a bacterial mechanosensitive channel into liposomal membranes. FASEB J. 2015 Oct;29(10):4334–45. doi:10.1096/fj.15-275198

62. Smart OS, Neduvelil JG, Wang X, Wallace BA, Sansom MSP. HOLE: A program for the analysis of the pore dimensions of ion channel structural models. J Mol Graph. 1996 Dec;14(6):354–60. doi:10.1016/S0263-7855(97)00009-X

63. Flegler VJ, Rasmussen A, Hedrich R, Rasmussen T, Böttcher B. Mechanosensitive channel engineering: A study on the mixing and matching of YnaI and MscS sensor paddles and pores. Nat Commun. 2025 Aug 23;16(1):7881. doi:10.1038/s41467-025-63253-0

64. Cox CD, Zhang Y, Zhou Z, Walz T, Martinac B. Cyclodextrins increase membrane tension and are universal activators of mechanosensitive channels. Proc Natl Acad Sci. 2021 Sep 7;118(36):e2104820118. doi:10.1073/pnas.2104820118

65. Hiotis G, Walz T. The bacterial mechanosensitive channel MscM gates through concerted changes in its transmembrane and cytoplasmic domains. Nat Commun. 2026 Jul 22. doi:10.1038/s41467-026-75798-9

66. Sukharev SI, Sigurdson WJ, Kung C, Sachs F. Energetic and Spatial Parameters for Gating of the Bacterial Large Conductance Mechanosensitive Channel, MscL. J Gen Physiol. 1999 Apr 1;113(4):525–40. doi:10.1085/jgp.113.4.525

67. Häse CC, Le Dain AC, Martinac B. Purification and Functional Reconstitution of the Recombinant Large Mechanosensitive Ion Channel (MscL) of Escherichia coli. J Biol Chem. 1995 Aug;270(31):18329–34. doi:10.1074/jbc.270.31.18329

68. Schumann U, Edwards MD, Li C, Booth IR. The conserved carboxy-terminus of the MscS mechanosensitive channel is not essential but increases stability and activity. FEBS Lett. 2004 Aug 13;572(1–3):233–7. doi:10.1016/j.febslet.2004.07.045

69. Abramson J, Adler J, Dunger J, Evans R, Green T, Pritzel A, et al. Accurate structure prediction of biomolecular interactions with AlphaFold 3. Nature. 2024 Jun 13;630(8016):493–500. doi:10.1038/s41586-024-07487-w

70. Becker M, Krämer R. MscCG from Corynebacterium glutamicum: functional significance of the C-terminal domain. Eur Biophys J. 2015 Oct;44(7):577–88. doi:10.1007/s00249-015-1041-x

71. Yim SS, An SJ, Kang M, Lee J, Jeong KJ. Isolation of fully synthetic promoters for high-level gene expression in *Corynebacterium glutamicum*. Biotechnol Bioeng. 2013 Nov;110(11):2959–69. doi:10.1002/bit.24954

72. Ohi M, Li Y, Cheng Y, Walz T. Negative staining and image classification — powerful tools in modern electron microscopy. Biol Proced Online. 2004 Jan;6(1):23–34. doi:10.1251/bpo70

73. Cheng A, Negro C, Bruhn JF, Rice WJ, Dallakyan S, Eng ET, et al. Leginon: New features and applications. Protein Sci. 2021 Jan;30(1):136–50. doi:10.1002/pro.3967

74. Mastronarde DN. Automated electron microscope tomography using robust prediction of specimen movements. J Struct Biol. 2005 Oct;152(1):36–51. doi:10.1016/j.jsb.2005.07.007

75. Meng EC, Goddard TD, Pettersen EF, Couch GS, Pearson ZJ, Morris JH, et al. UCSF CHIMERAX : Tools for structure building and analysis. Protein Sci. 2023 Nov;32(11):e4792. doi:10.1002/pro.4792

76. Emsley P, Lohkamp B, Scott WG, Cowtan K. Features and development of *Coot*. Acta Crystallogr D Biol Crystallogr. 2010 Apr 1;66(4):486–501. doi:10.1107/S0907444910007493

77. Afonine PV, Poon BK, Read RJ, Sobolev OV, Terwilliger TC, Urzhumtsev A, et al. Real-space refinement in *PHENIX* for cryo-EM and crystallography. Acta Crystallogr Sect Struct Biol. 2018 Jun 1;74(6):531–44. doi:10.1107/S2059798318006551

78. Sanchez-Garcia R, Gomez-Blanco J, Cuervo A, Carazo JM, Sorzano COS, Vargas J. DeepEMhancer: a deep learning solution for cryo-EM volume post-processing. Commun Biol. 2021 Jul 15;4(1):874. doi:10.1038/s42003-021-02399-1

79. Jo S, Kim T, Iyer VG, Im W. CHARMM-GUI: A web-based graphical user interface for CHARMM. J Comput Chem. 2008 Aug;29(11):1859–65. doi:10.1002/jcc.20945

80. Lee J, Cheng X, Swails JM, Yeom MS, Eastman PK, Lemkul JA, et al. CHARMM-GUI Input Generator for NAMD, GROMACS, AMBER, OpenMM, and CHARMM/OpenMM Simulations Using the CHARMM36 Additive Force Field. J Chem Theory Comput. 2016 Jan 12;12(1):405–13. doi:10.1021/acs.jctc.5b00935

81. Durell SR, Brooks BR, Ben-Naim A. Solvent-Induced Forces between Two Hydrophilic Groups. J Phys Chem. 1994 Feb 1;98(8):2198–202. doi:10.1021/j100059a038

82. Jorgensen WL, Chandrasekhar J, Madura JD, Impey RW, Klein ML. Comparison of simple potential functions for simulating liquid water. J Chem Phys. 1983 Jul 15;79(2):926–35. doi:10.1063/1.445869

83. Steinbach PJ, Brooks BR. New spherical-cutoff methods for long-range forces in macromolecular simulation. J Comput Chem. 1994 Jul;15(7):667–83. doi:10.1002/jcc.540150702

84. Darden T, York D, Pedersen L. Particle mesh Ewald: An *N* ⋅log( *N* ) method for Ewald sums in large systems. J Chem Phys. 1993 Jun 15;98(12):10089–92. doi:10.1063/1.464397

85. Essmann U, Perera L, Berkowitz ML, Darden T, Lee H, Pedersen LG. A smooth particle mesh Ewald method. J Chem Phys. 1995 Nov 15;103(19):8577–93. doi:10.1063/1.470117

86. Bernetti M, Bussi G. Pressure control using stochastic cell rescaling. J Chem Phys. 2020 Sep 21;153(11):114107. doi:10.1063/5.0020514

87. Bussi G, Donadio D, Parrinello M. Canonical sampling through velocity rescaling. J Chem Phys. 2007 Jan 7;126(1):014101. doi:10.1063/1.2408420

88. Hess B. P-LINCS: A Parallel Linear Constraint Solver for Molecular Simulation. J Chem Theory Comput. 2008 Jan 1;4(1):116–22. doi:10.1021/ct700200b

89. Abraham MJ, Murtola T, Schulz R, Páll S, Smith JC, Hess B, et al. GROMACS: High performance molecular simulations through multi-level parallelism from laptops to supercomputers. SoftwareX. 2015 Sep;1–2:19–25. doi:10.1016/j.softx.2015.06.001

90. Huang J, Rauscher S, Nawrocki G, Ran T, Feig M, De Groot BL, et al. CHARMM36m: an improved force field for folded and intrinsically disordered proteins. Nat Methods. 2017 Jan;14(1):71–3. doi:10.1038/nmeth.4067

91. Huang J, MacKerell AD. CHARMM36 all-atom additive protein force field: Validation based on comparison to NMR data. J Comput Chem. 2013 Sep 30;34(25):2135–45. doi:10.1002/jcc.23354

92. Roux B. The Membrane Potential and its Representation by a Constant Electric Field in Computer Simulations. Biophys J. 2008 Nov;95(9):4205–16. doi:10.1529/biophysj.108.136499

93. Michaud-Agrawal N, Denning EJ, Woolf TB, Beckstein O. MDAnalysis: A toolkit for the analysis of molecular dynamics simulations. J Comput Chem. 2011 Jul 30;32(10):2319–27. doi:10.1002/jcc.21787

94. Gowers R, Linke M, Barnoud J, Reddy T, Melo M, Seyler S, et al. MDAnalysis: A Python Package for the Rapid Analysis of Molecular Dynamics Simulations. In. Austin, Texas; 2016 [cited 2026 Jun 17]. p. 98–105. Available from: https://doi.curvenote.com/10.25080/Majora-629e541a-00e doi:10.25080/Majora-629e541a-00e

95. Katoh K, Standley DM. MAFFT Multiple Sequence Alignment Software Version 7: Improvements in Performance and Usability. Mol Biol Evol. 2013 Apr 1;30(4):772–80. doi:10.1093/molbev/mst010

96. Capella-Gutiérrez S, Silla-Martínez JM, Gabaldón T. trimAl: a tool for automated alignment trimming in large-scale phylogenetic analyses. Bioinformatics. 2009 Aug 1;25(15):1972–3. doi:10.1093/bioinformatics/btp348

97. Minh BQ, Schmidt HA, Chernomor O, Schrempf D, Woodhams MD, Von Haeseler A, et al. IQ-TREE 2: New Models and Efficient Methods for Phylogenetic Inference in the Genomic Era. Teeling E, editor. Mol Biol Evol. 2020 May 1;37(5):1530–4. doi:10.1093/molbev/msaa015

98. Letunic I, Bork P. Interactive Tree of Life (iTOL) v6: recent updates to the phylogenetic tree display and annotation tool. Nucleic Acids Res. 2024 Jul 5;52(W1):W78–82. doi:10.1093/nar/gkae268

99. Waterhouse AM, Procter JB, Martin DMA, Clamp M, Barton GJ. Jalview Version 2—a multiple sequence alignment editor and analysis workbench. Bioinformatics. 2009 May 1;25(9):1189–91. doi:10.1093/bioinformatics/btp033

100. Crooks GE, Hon G, Chandonia JM, Brenner SE. WebLogo: A Sequence Logo Generator: Figure 1. Genome Res. 2004 Jun;14(6):1188–90. doi:10.1101/gr.849004

