## Supplementary material for "The MscS channel from *Corynebacterium glutamicum* uses a non-canonical gating mechanism": CgMscS ms Nat Comm.pdf

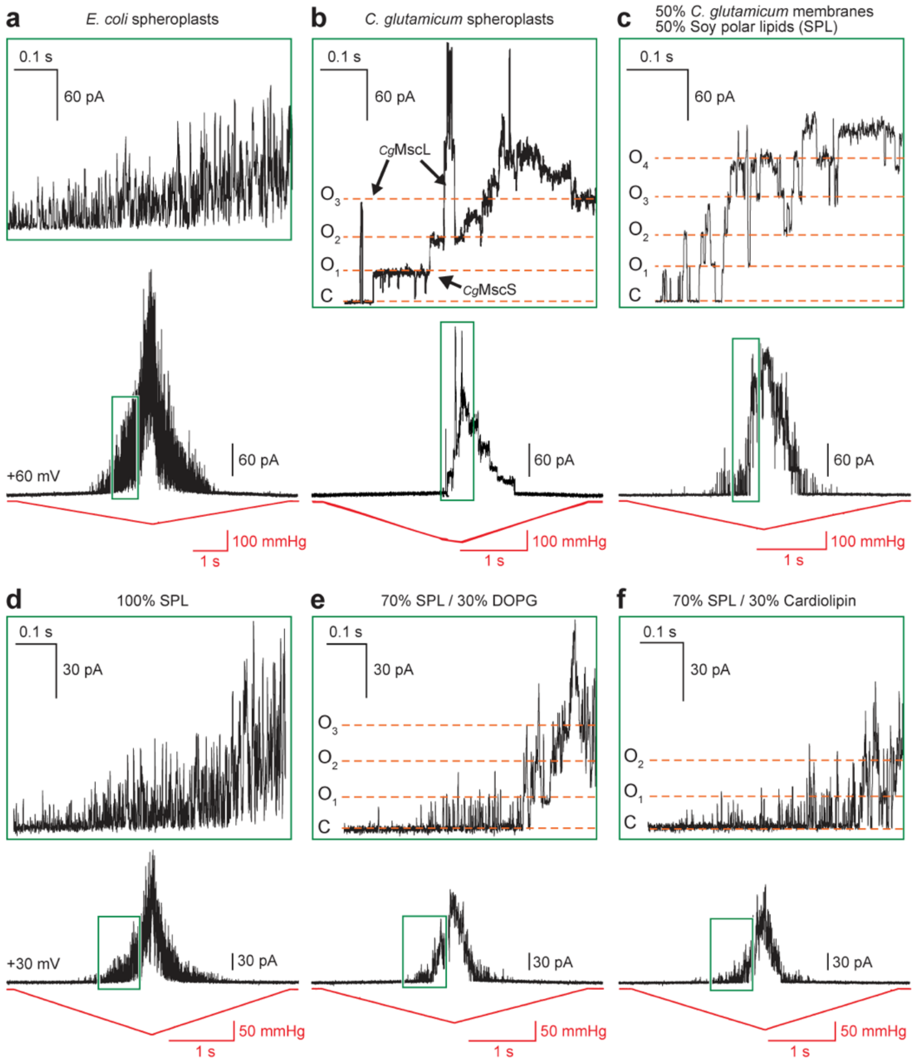


**Fig. 1 | Anionic lipids stabilize the open state of *Cg*MscS. a-c** Representative traces of currents in response to ramp-pressure stimuli recorded from giant spheroplasts prepared from *E. coli* expressing *Cg*MscS (a), giant spheroplasts prepared from *C. glutamicum* expressing *Cg*MscS (adapted from Nakayama *et al.*, *Sci Rep* 2018) (b), and fusion vesicles obtained by fusing membrane vesicles prepared from *Cg*MscS-expressing *C. glutamicum* with soy polar lipids (SPL) liposomes (c). **d-f** Representative traces of currents in response to ramp-pressure stimuli recorded from *Cg*MscS proteoliposomes reconstituted with 100% SPL (d), 70% SPL and 30% DOPG (e), and 70% SPL and 30% cardiolipin (f). The top panels are zoomed-in views of the regions indicated by the boxes in the bottom panels to show single-channel current levels. Red line: pressure; black line: current; horizontal orange dashed lines: levels when all *Cg*MscS channels are closed (C) and when one, two, three or four *Cg*MscS channels are open (O1 – O4).

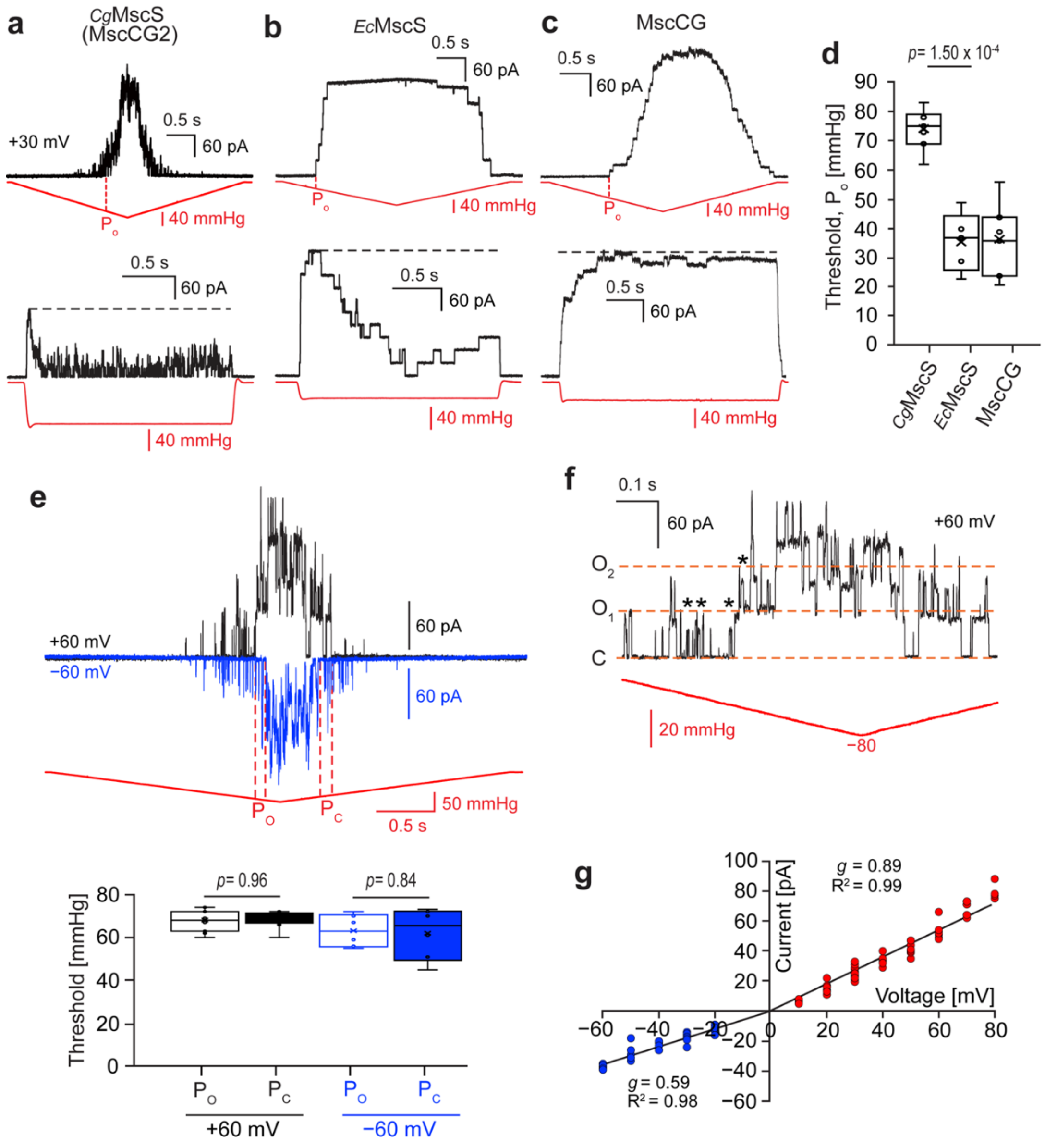


**Fig. 2 | Channel characteristics of *Cg*MscS**. **a-c** Representative traces of currents in response to ramp (top panels) and step (bottom panels) pressure stimuli. Panel a shows current traces recorded from fusion vesicles formed from *Cg*MscS-containing *C. glutamicum* membrane vesicles and SPL liposomes (n = 21). The step-pressure recording shows rapid *Cg*MscS activation followed by current decay during sustained pressure. Panel b shows current traces recorded from fusion vesicles formed from *Ec*MscS-containing *E. coli* membrane vesicles and SPL liposomes (n = 5). Panel c shows current traces recorded from fusion vesicles formed from MscCG-containing *C. glutamicum* membrane vesicles and SPL liposomes (n = 7). The step-pressure recording shows slower activation and sustained currents with little or no inactivation. Red line: pressure; black line: current; $P_o$ in top panels: first full channel opening; horizontal black dashed lines in bottom panels: peak current levels. **d** Comparison of activation thresholds, measured as the pressure of the first full channel opening, for *Cg*MscS (n = 7), *Ec*MscS (n = 5) and MscCG (n = 7). Statistical analysis: two-sided Welch's unequal-variance *t*-test. Box plot shows median (centre line), interquartile range (box) and full data range (whiskers). **e** Top panel: Representative traces of currents in response to ramp-pressure stimuli recorded from *Cg*MscS-containing fusion vesicles at voltages of +60 mV and −60 mV (n = 6). Red line: pressure; black/blue line: current; $P_o$: first full channel opening; $P_c$: last channel closing. Bottom panel: The opening and closing thresholds show no statistically significant differences, indicating that the channels have no gating hysteresis at both positive and negative voltages. Statistical analysis: two-sided Welch's unequal-variance *t*-test. Box plot shows median (centre line), interquartile range (box) and full data range (whiskers). **f** Single-channel conductance analysis of WT *Cg*MscS. Currents were recorded from *Cg*MscS-containing fusion vesicles at +60 mV (n = 11). Red line: pressure; black line: current; horizontal orange dashed lines: levels when all *Cg*MscS channels are closed (C) and when one or two *Cg*MscS channels are open (O1, O2), asterisks: subconductive levels. **g** Current–voltage relationship for *Cg*MscS. Currents were recorded from *Cg*MscS-containing fusion vesicles at voltages ranging from −60 mV to +80 mV (more than three independent patch membranes per voltage). The slope conductances (*g*) and coefficients of determination ($R^2$) were calculated from independent linear fits to the measurements obtained with positive (red) and negative (blue) voltages.

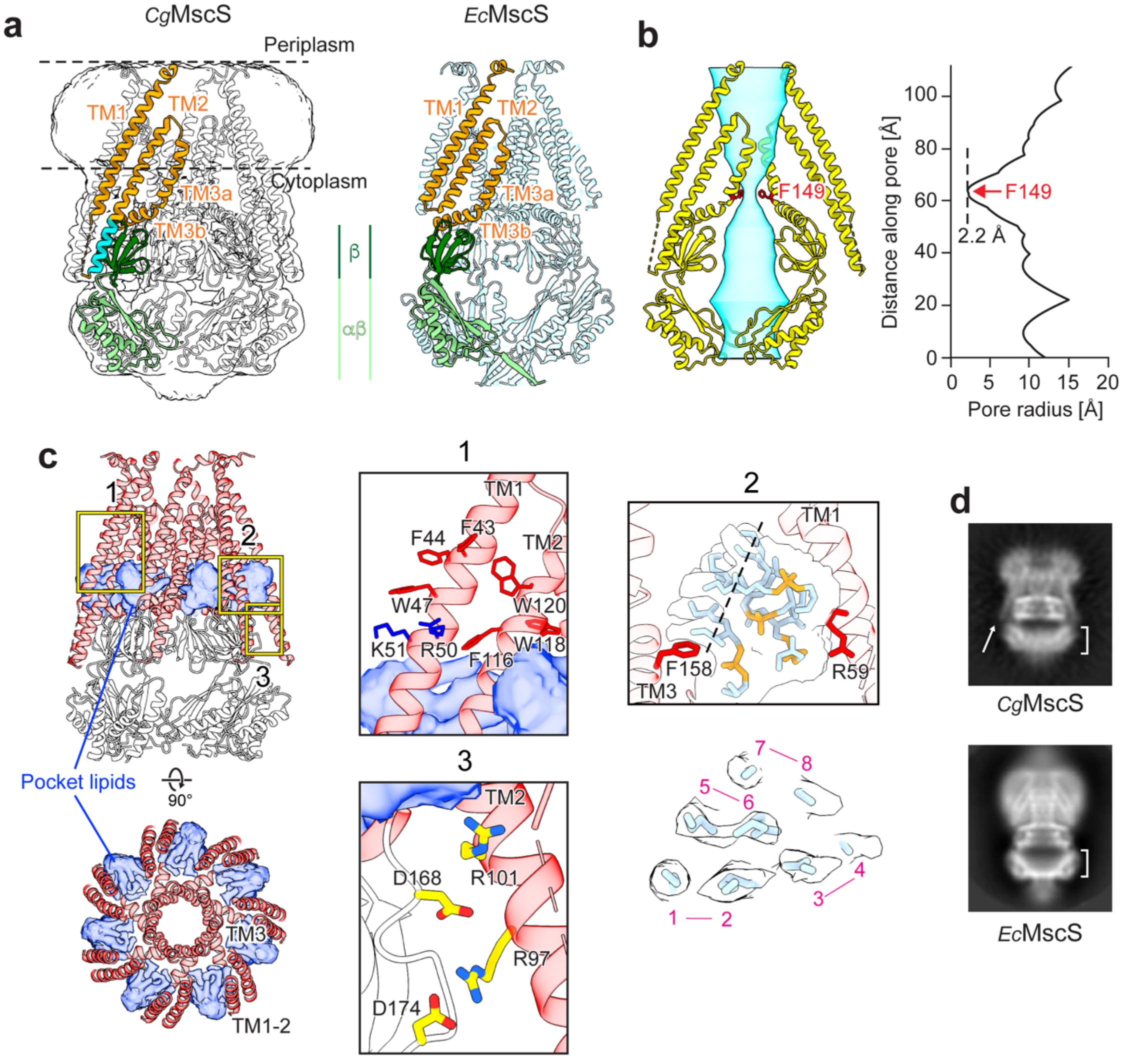


**Fig. 3 | Cryo-EM structure of WT *Cg*MscS in DOPG nanodiscs**. **a** Cryo-EM structures of *Cg*MscS in DOPG nanodiscs (left) and *Ec*MscS in DOPC nanodics (right; PDB: 6VYK) (22). The *Cg*MscS density map is shown as transparent white surface, and the models for the *Cg*MscS and *Ec*MscS heptamers are shown in white and light blue, respectively. One protomer in each channel is color-coded: orange: TM1-2 (sensor paddle), TM3a (pore-lining helix) and TM3b; green: cytoplasmic cage (β subdomain in dark green; αβ subdomain in light green); cyan: TM2 extension unique to *Cg*MscS. The horizontal dashed lines indicate the membrane boundary predicted by the Orientations of Proteins in Membranes database (https://opm.phar.umich.edu). **b** The volume representation (left panel) and linear profile of the pore radius (right panel) generated with HOLE (62) show that the pore in *Cg*MscS is closed and that the constriction site is formed by the aromatic

Phe149 residue. The vertical dashed line shows the pore radius at the constriction site. **c** Views perpendicular (top panel) and parallel to the membrane (bottom panel) of the *Cg*MscS heptamer shown in red (TM helices) and white (cytoplasmic cage) ribbon representation. Densities occupying the hydrophobic extramembranous cavities that likely represent pocket lipids are shown as blue transparent surfaces. Box 1 shows a zoomed-in view of aromatic residues (Trp and Phe; red) and positively charged residues (Arg and Lys; blue) near the cytoplasmic membrane boundary that may anchor the position of TM1 and TM2. Box 2 shows a zoomed-in view of the four lipids modeled into the pocket density, and the panel below shows a section through the pocket density at the position indicated in Box 2, revealing density for the eight acyl chains. Box 3 shows a zoomed-in view of the putative electrostatic interactions between arginine residues in the TM2 extension and aspartate residues in the β subdomain that link the sensor paddle to the cytoplasmic cage. **d** 2D-class averages showing side views of *Cg*MscS (top panel) and *Ec*MscS (bottom panel, from Zhang et al., Nature, 2022). The arrow indicates the extended TM2 in *Cg*MscS and the brackets show the αβ subdomain that is well-defined in *Ec*MscS but smeared out in *Cg*MscS.

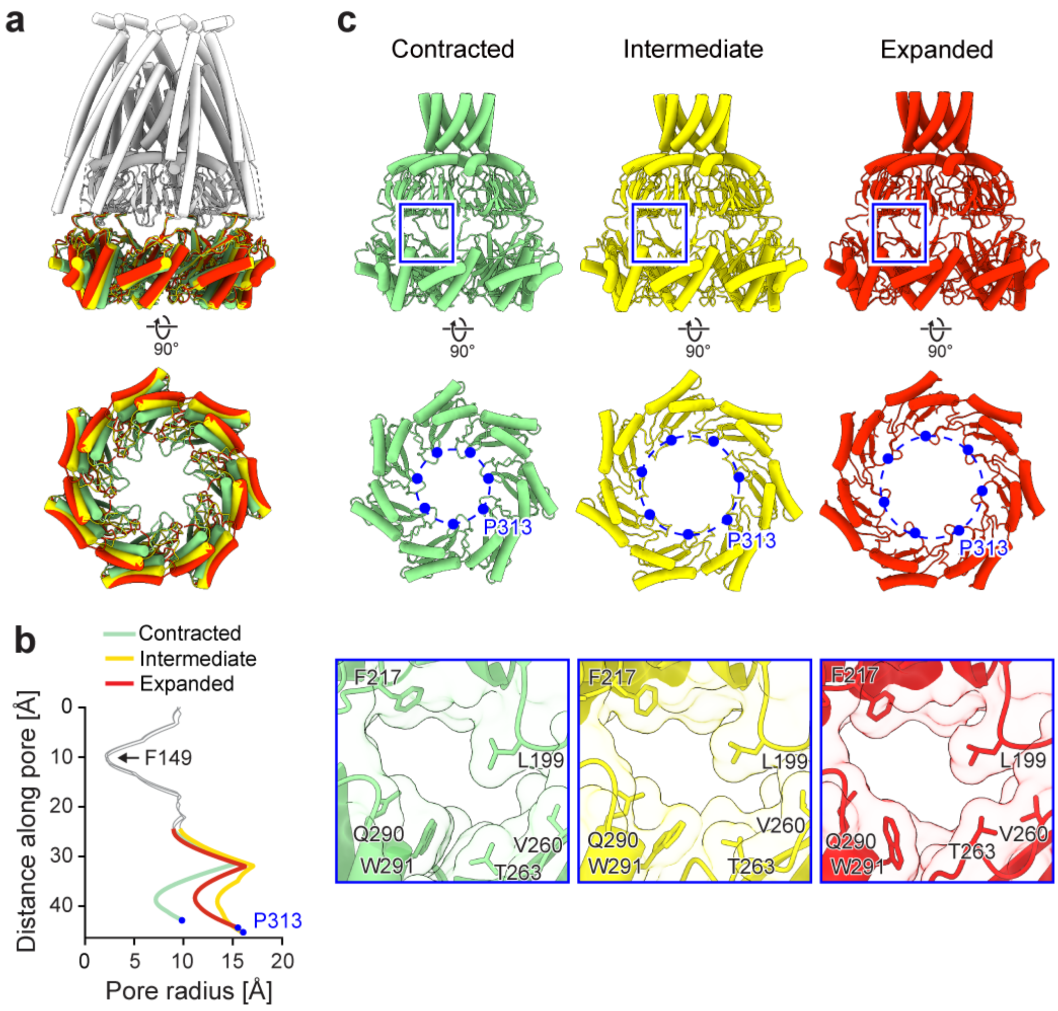


**Fig. 4 | Expansion/contraction of the αβ subdomain of WT *Cg*MscS.** **a** Overlay of the most contracted (light green), intermediate (yellow), and most expanded (red) conformations of the αβ subdomain. Views perpendicular (top panels) and parallel to the membrane (bottom panels) are shown. **b** HOLE radius profiles in the region of TM3, the β subdomain, and the αβ subdomain (residues 134–313) of *Cg*MscS with the αβ subdomain in the three different conformations. The portions of the profiles corresponding to TM3 and the β subdomain are shown in gray, whereas those corresponding to the αβ subdomain in the most contracted, intermediate and most expanded conformation are shown in light green, yellow and red, respectively. The constriction at Phe149, indicated by the arrow, is essentially unchanged between the three conformations. The blue filled circles mark the pore radius at the level of the last modeled residue, Pro313, illustrating that the radius of the axial opening in the cage is smaller in the most contracted conformation. **c** Comparison of the αβ subdomain and the lateral fenestrations in the most contracted (light green), intermediate (yellow), and most expanded (red) conformations. Views are shown parallel to the membrane for the cytoplasmic cage and TM3 (top panels) and perpendicular to the membrane from the cytoplasmic side for the αβ subdomain (middle panels). The blue filled circles mark the positions of Pro313, and the blue dashed circles indicate the diameter of the ring formed by the Pro313 residues. The bottom panels show close-up views of the lateral fenestrations indicated by the boxes in the top panels, illustrating that the openings are similar.

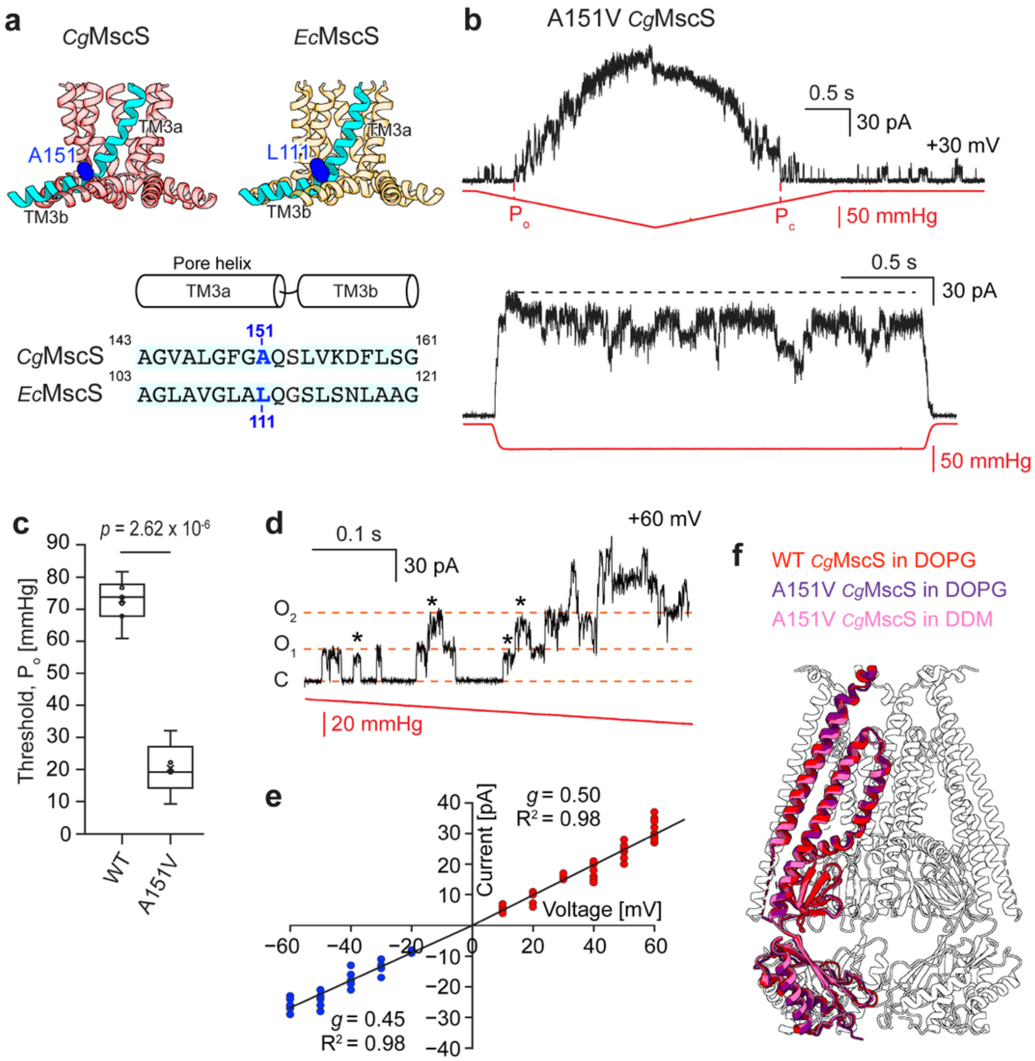


**Fig. 5 | Functional and structural characterization of A151V *Cg*MscS. a** Position of the A151V mutation in the kink region of the pore-lining TM3a helix of *Cg*MscS (transparent red, top left panel) and the corresponding residue L111 in *Ec*MscS (transparent orange, top right panel). One TM3 helix is shown in cyan, and residues A151 in *Cg*MscS and L111 in *Ec*MscS are shown as blue spheres. The bottom panel shows a sequence alignment of the TM3 segments surrounding the kink region in *Cg*MscS and *Ec*MscS, with the corresponding residues highlighted in blue. **b** Patch-clamp recordings of A151V *Cg*MscS in fusion vesicles in response to ramp (top panel) and step (bottom panel) pressure stimuli (n = 18, each). Red line: pressure; black line: current; $P_o$: first full channel opening; $P_c$: last channel closing; horizontal black dashed line: peak current level. **c** Comparison of activation thresholds, measured as the pressure of the first full channel opening, for WT *Cg*MscS (n = 7) and A151V *Cg*MscS (n = 5). Statistical analysis: two-sided Welch's unequal-variance t-test. Box plot shows median (center line), interquartile range (box), and full data range (whiskers). **d** Single-channel conductance analysis of A151V *Cg*MscS. Currents were recorded from *Cg*MscS-containing fusion vesicles at +60 mV (n = 18). Red line: pressure; black line: current; horizontal orange dashed lines: levels when all *Cg*MscS channels are closed (C) and when one or two *Cg*MscS channels are open (O1, O2), asterisks: subconductive levels. **e** Current–voltage relationship for A151V *Cg*MscS. Currents were recorded from A151V *Cg*MscS-containing fusion vesicles at voltages ranging from −60 mV to +60 mV (more than three independent patch membranes per voltage). The slope conductances (*g*) and coefficients of determination ($R^2$) were calculated from independent linear fits to the measurements obtained with positive (red) and negative (blue) voltages. **f** Overlay of the cryo-EM structures of WT *Cg*MscS in DOPG nanodiscs (red), A151V *Cg*MscS in DOPG nanodiscs (purple), and A151V *Cg*MscS in DDM (pink), showing that the A151V mutant remains in a closed conformation.

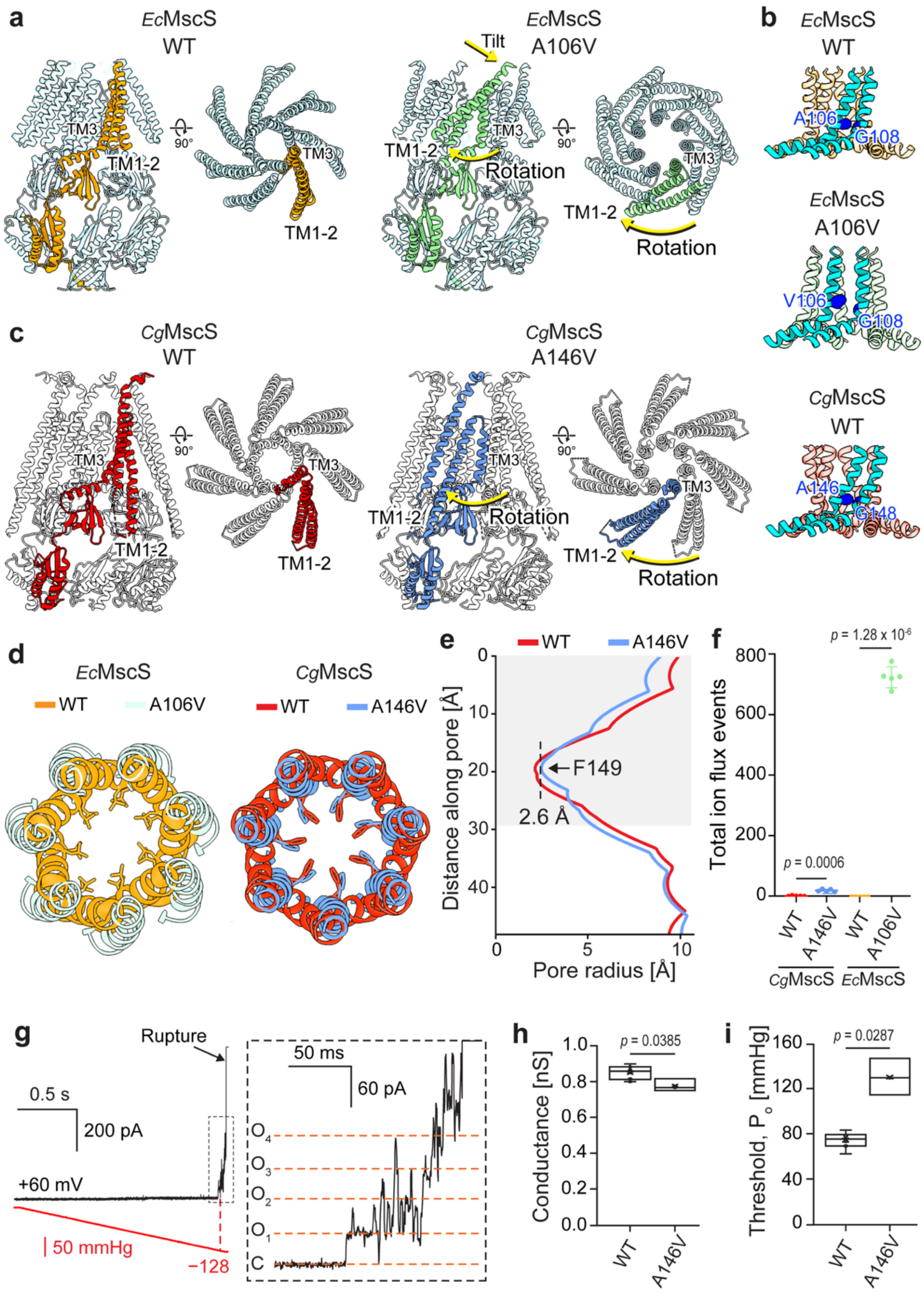


**Fig. 6 | Structural and functional characterization of A146V *Cg*MscS. a** Cryo-EM structures of WT *Ec*MscS in the closed conformation (left; PDB: 2OAU; one protomer in orange) (14) and A106V *Ec*MscS in open conformation (right; PDB: 2VV5; one protomer in green) (18), showing that the TM1-2 paddle rotates and tilts. **b** Intersubunit packing interactions between adjacent TM3a helices in WT *Ec*MscS in the closed state (top), A106V *Ec*MscS in the open state (middle), and WT *Cg*MscS in the closed state (bottom). Two neighboring protomers are shown in cyan, and glycine and alanine residues involved in helical packing interactions are shown as blue spheres. **c** Cryo-EM structures of WT *Cg*MscS in the closed conformation (left, one protomer in red) and A146V *Cg*MscS in the inactivated conformation (right; one protomer in blue), showing that the TM1-2 paddle only undergoes a rotation but no tilt. **d** Arrangement of the TM3a helices in closed WT *Ec*MscS (orange) and open A106V *Ec*MscS (light green) (left), and in closed WT *Cg*MscS (red) and inactivated A146V *Cg*MscS (blue) (right). The side chains of Leu105 and Leu109 in *Ec*MscS and of Phe149 in *Cg*MscS, which form the pore constrictions, are shown. **e** Pore radius profiles generated with HOLE for WT (red) and A146V *Cg*MscS (blue). The light gray box indicates the pore region lined by the TM3a helices. **f** Total ion-flux events observed in 200-ns molecular-dynamics simulations for WT and mutant *Ec*MscS and *Cg*MscS. Data plotted as mean ± SD (n = 5). Statistical analysis: two-sided Welch's unequal-variance *t*-test. **g** Currents were recorded from A146V *Cg*MscS-containing fusion vesicles in response to increasing negative pressure (n = 3). Red line: pressure; black line: current; vertical red dashed line: first full channel opening. The right panel shows a zoomed-in view of the region indicated by a box in the left panel to resolve single-channel conductance levels. Horizontal dashed lines: levels when all channels are closed (C) and when one, two, three or four channels are open (O1 – O4). **h, i** Comparison of channel conductance (h) for WT (n = 10) and A146V *Cg*MscS (n = 3) and of activation thresholds, measured as the pressure of the first full channel opening (i), for WT (n = 7) and A146V *Cg*MscS (n = 3). Statistical analysis: two-sided Welch's unequal-variance t-test. Box plots show median (center line), interquartile range (box), and full data range (whiskers).

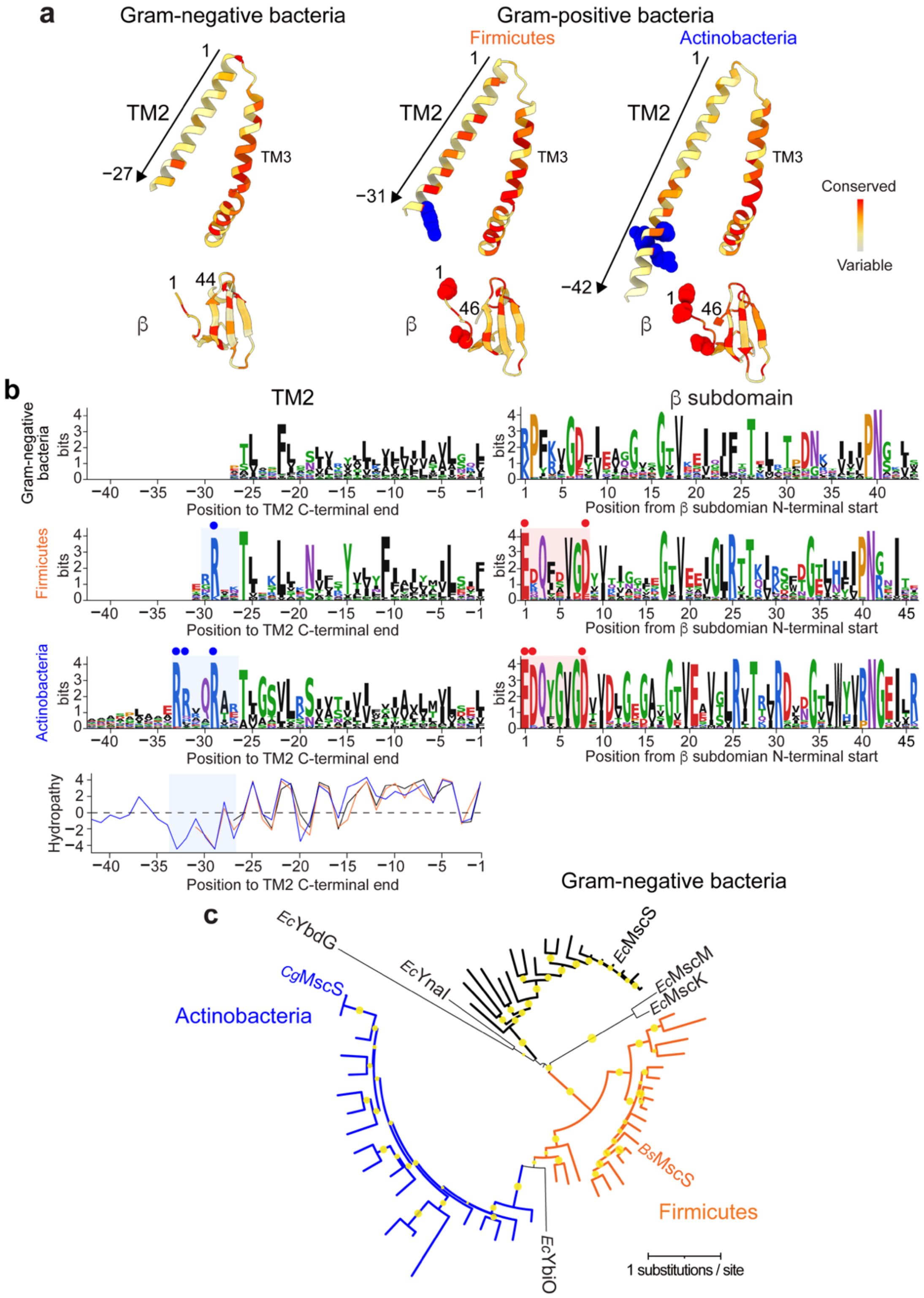


**Fig. 7 | Conservation and phylogenetic relationships of MscS homologs from Gram-negative and Gram-positive bacteria. a** Sequence conservation mapped onto the structures of representative homologs from each bacterial phylum: *Ec*MscS for Gram-negative bacteria, *Bs*MscS for Firmicutes, and *Cg*MscS for Actinobacteria. Conservation is shown as a gradient from highly variable (light gray) to highly conserved (red). Conserved arginine residues in TM2 are shown as blue spheres, and conserved acidic residues in the β subdomain are shown as red spheres. **b** Sequence-logo analyses of TM2 (left panels) and the β subdomain (right panels) of MscS homologs from Gram-negative bacteria, Firmicutes, and Actinobacteria. Logos were generated from alignments of 2,758 homologs from Gram-negative bacteria, 1,101 homologs from Firmicutes, and 817 homologs from Actinobacteria. Blue circles indicate conserved arginine residues in the cytoplasmic extension of TM2, and red circles indicate conserved acidic residues in the β subdomain. Hydropathy profiles of the consensus TM2 sequences are shown in the bottom panel. **c** Maximum-likelihood phylogenetic tree of representative MscS homologs from Gram-negative bacteria, Firmicutes, and Actinobacteria. Branches corresponding to homologs from Gram-negative bacteria, Firmicutes, and Actinobacteria are shown in black, orange, and blue, respectively. The scale bar indicates the number of amino acid substitutions per site. Circles at internal nodes indicate bootstrap support values greater than 60%.

**FUNDING STATEMENT**

This work was supported by National Institutes of Health grant R01 GM144581 (T.W.) and the Stavros Niarchos Foundation (SNF) as part of its grant to the SNF Institute for Global Infectious Disease Research at The Rockefeller University (Y.N., T.W.).

**Author contributions**

Y.N. and T.W. conceived the study. T.W. supervised the research. Y.N. performed the cryo-EM and patch-clamp experiments and the sequence conservation and phylogenetic analyses. G.H. performed the MD simulations. Y.N. and G.H. analyzed the results. Y.N. prepared the manuscript with assistance of T.W. All authors reviewed and agreed with the manuscript.

**Competing interests**

The authors declare no competing interests.
